# Nonstop nanometric resolution of randomly moving point scatterers with focused light

**DOI:** 10.64898/2026.02.09.704780

**Authors:** Thomas A. Hensel, Ole L. Schwarz, Tim Karrasch, Kerstin Göpfrich, Stefan W. Hell

**Author notes:** Correspondence: Stefan W. Hell.

## Abstract

In established super-resolution fluorescence microscopy, resolving multiple fluorescent molecules at sub-diffraction distances requires the molecules to emit sequentially so that they become discernible from their neighbors one after another. Simultaneous tracking of multiple fluorophores that are only a few nanometers apart is thus conceptually and practically impossible. We have recently shown that probing a sub-diffraction region with an excitation beam featuring an intensity zero, i.e., MINFLUX, super-resolves and tracks closely packed fluorophores without interruption. Here, we provide a conceptual framework for resolving and tracking constantly emitting fluorophores – more generally, point scatterers – that undergo random changes in position. In particular, we show that the detection rates available in fluorescence microscopy are sufficient to track sub-10 nm distance changes within micro-to milliseconds. By using a DNA origami construct with a fixed and a movable fluorophore as a proxy, we prove the concept that thermally driven conformational changes of biomolecules are continuously detectable with visible light. Conformational changes of the DNA nanostructure leading to random jumps in distance of about 10 nm between two labels are registered within about a millisecond. Our work paves the way towards super-resolving complex conformational transitions of individual biomolecules with focused light.

## Introduction

Since its first demonstration, super-resolution fluorescence microscopy or nanoscopy has evolved into a powerful method for mapping the distribution of fluorophore-labeled biomolecules in cells. By providing (sub-)nanometer localization accuracy of single fluorophores, some of the latest super-resolution methods, such as those called MINFLUX (1) and MINSTED (2) have become increasingly suitable for studying the biomolecules themselves. Particularly MINFLUX opens up the possibility to follow conformational changes of individual proteins just by measuring the positional change of two or more identical fluorophore tags bound to the protein. In fact, it has recently been shown that MINFLUX can accurately classify protein conformations by measuring differences in distance between two photoactivatable fluorophore tags (3). Nonetheless, all established super-resolution methods, including the reported implementation of MINFLUX, cannot observe the conformational rearrangement in action, and hence not its mechanistic details.

The reason is that the observation of the repositioning of several fluorophore tags requires uninterrupted monitoring and resolution of all tags at the same time, which is not compatible with the ON/OFF working principle of established super-resolution microscopy. In the case of individual fluorophores, the ON/OFF principle clearly demands that only a single fluorophore is able to emit at any point in time. As the monitoring of collective positional changes of the fluorophores is thus fundamentally impossible, direct observation of conformational changes with established super-resolution methods is very limited or simply not viable. Detecting just a single moving fluorophore at a time is not sufficient, because the movement of a single fluorophore tag can only indicate the result of a conformational change, such as a translation in the case of motor proteins (4, 5), or a trivial movement of the protein as a whole. Generally speaking, the ON/OFF separation principle does not allow the observation of collective fluorophore movements that are faster than the time span required for all the fluorophores to sequentially undergo the ON/OFF cycle. As the protein conformational change and most other biomolecular processes occur on much shorter time spans, the current super-resolution paradigm is not suitable for observing such changes on fundamental grounds.

To overcome this limitation, we recently introduced a sub-diffraction resolution principle that does not require a transient state transition for separation (6). Concretely, we showed that the MINFLUX principle of probing the object with an illumination intensity minimum resolves a countable number of point scatterers at distances of a tiny fraction of the illumination wavelength – even if they all produce signal at the same time. Since an ON/OFF state transition, such as switching or uncaging, is not required, the position of all the scatterers can be interrogated without interruption. Thus, MINFLUX opened up a new avenue towards direct observation of conformational changes of single proteins.

In the report introducing this new principle (6), we also resolved two fluorophores that changed their position in the x,y-focal plane of the microscope. However, those fluorophores were tightly bound to a glass cover slip and their movement was artificially imparted by the microscope stage. Hence, the fluorophores could neither change their distance (pre-set either at 15 nm or 30 nm) nor their mutual orientation, which is in stark contrast to a real biomolecule where the positions of the fluorophores change arbitrarily.

Here, we show that MINFLUX resolves two constantly emitting identical fluorophores non-stop, as they randomly change their Euclidian distance at the nanometer scale through thermal motion. The rapid distance change was realized by attaching one of the fluorophores to a movable DNA duplex lever arm anchored on a DNA origami sheet on a glass cover slip. The second fluorophore was firmly placed on the sheet itself. The origami also featured two transient binding sites for a complementary single-stranded DNA overhang on the arm, one at a shorter and one at a longer distance to the fixed fluorophore, offering two well-defined inter-fluorophore distances upon binding. Like the DNA constructs used by Eilers et al. (7), for demonstrating fast tracking of single fluorophores by MINFLUX, the present DNA nanostructure allowed us to demonstrate that the MINFLUX-typical spatial probing with an intensity minimum of excitation light resolves on the (sub)millisecond time scale two identical fluorophores undergoing random nanometric distance changes.

## Results

Since the position of the fluorophores is regarded as a proxy of the conformational state of a biomolecule, we now define the set of positions of the two fluorophores as the positional state of the two-fluorophore system. Thus, a biomolecular conformation is tightly linked to a positional state of the fluorophores. In most cases, the quantification of the latter can be broken down to measuring i) the changes of their distance and orientation in space and ii) the changes of the center of mass (COM) of the fluorophores. Therefore, we first explore how these changes can be detected by probing with an illumination intensity minimum.

### Quantifying the change in distance between two scatterers

Consider an excitation beam with a line-shaped minimum (Fig. 1A) as generated by two interfering laser beams in the focal plane of a microscope (Fig. 1B). The contrast of the interference pattern resulting from potentially imperfect interference, is denoted with *ν*_0_ = (*I*_*max*_ − *I*_*min*_)/(*I*_*max*_ + *I*_*min*_), with *I*_*max*_ and *I*_*min*_ denoting the intensity at the interference maximum and minimum, respectively. The x- and y-orientation of the line-shaped minimum is readily interchanged by rotating the interfering beams in the pupil plane of the objective lens by 90°. Varying the phase difference Δ*φ* between the beams moves the minimum over a spatial interval *L* in the focal plane (Fig. 1C). Scanning this minimum across an ensemble of fluorescent molecules with varying orientations encodes the spatial distribution of the fluorophores in the deviation of the minimum of the collected fluorescence intensity from zero, i.e., in the minimum “depth”. Changes in fluorophore separation change this depth (Fig. 1D), implying that in return, changes in depth reveal changes in distance.

**Fig. 1.**
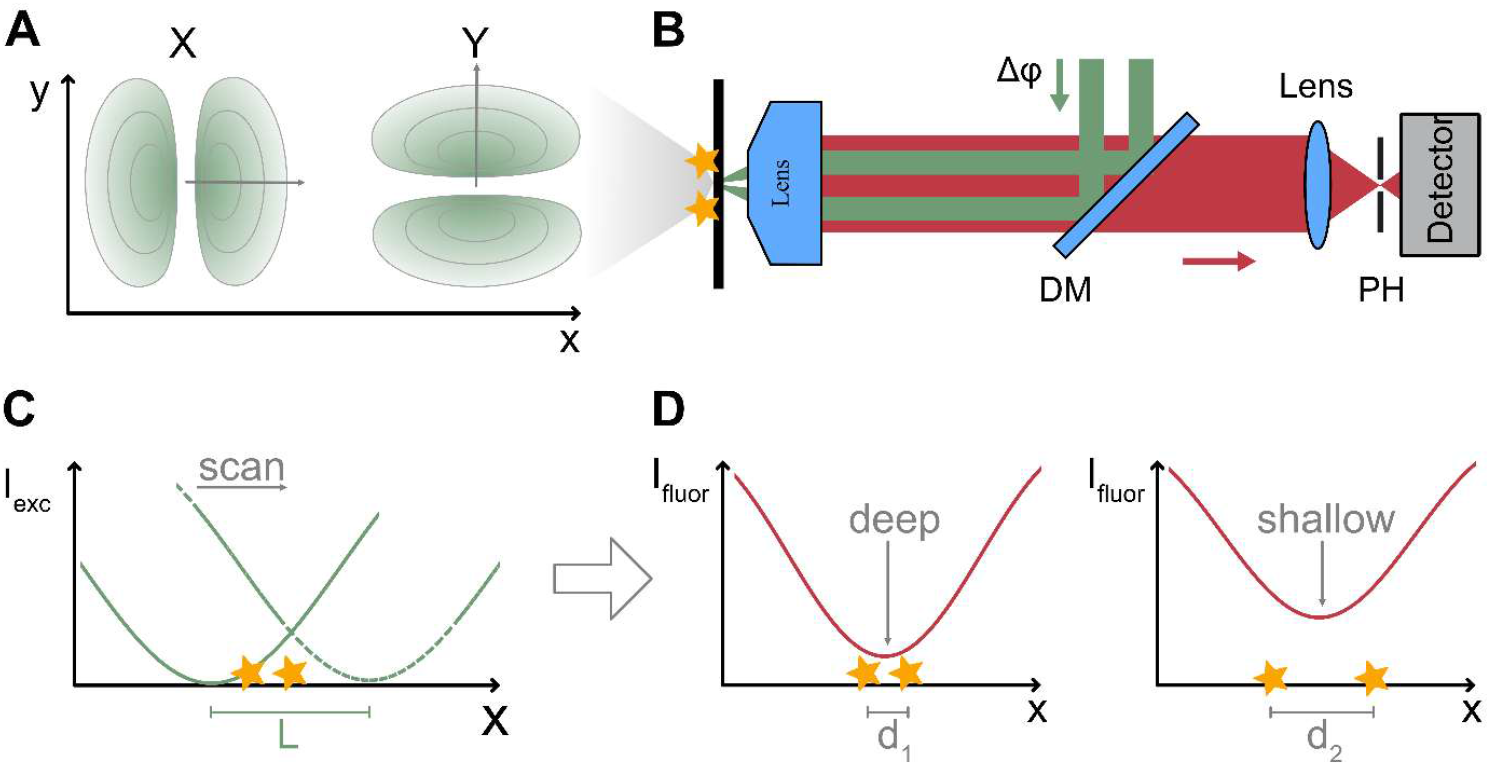
Nonstop separation and tracking of two or more constantly scattering, identical point objects by MINFLUX relies on scanning with an illumination beam featuring an intensity minimum. (**A**) A y- and x-oriented line-shaped minimum is scanned over the point objects (stars), to identify their positions in the x- and y-direction, respectively. (**B**) MINFLUX microscopy setup with a line-shaped minimum in the focal plane, produced by interfering two coherent laser beams (green) destructively at the focal point of the objective lens. Ramping the phase difference Δ*φ* between the two beams scans the minimum across a sub-diffraction range (< 75 nm). If fluorophores are used as point scatterers, the emitted fluorescence light (red) is collected by the lens and is registered, after passing a dichroic mirror (DM), by a detector placed behind a confocal pinhole (PH). (**C**) Scanning the y- or x-oriented minimum of the excitation beam *I*_*exc*_ along the x- or y-axis, respectively, over a distance L. (**D**) Since both the depth of the minimum of the fluorescence intensity profile *I*_*fluor*_ and its slope depend on the distance *d*_l,2_ between the scatterers, changes in distance Δ*d* = *d*_2_ − *d*_1_ are readily extractable from changes in depth and slope. The concept can be extended in three-dimensions and to several identical point scatterers of known number.

When probing two incoherent point scatterers, such as fluorophores, with a diffraction minimum, the co-localization precision scales linearly with the distance between the scatterers. In other words, contrary to probing with a diffraction maximum, the precision of the simultaneous localization of the fluorophores improves with the distances becoming smaller. Based on this promising scaling, we now derive the spatio-temporal resolution when jointly tracking two incoherent scatterers at a given scattering or fluorescence rate.

Consider two incoherent point scatterers with changing distance *d*, expressed in units of the standing-wave spatial period *λ*/2 of the illumination light. We now seek to establish the smallest discernable change in *d*. This problem can be rephrased as the ability to discriminate between two constellations of fluorophores with differing distances *d*_1_ and *d*_2_. Defining Δ*d* = |*d*_2_ − *d*_1_ | and the mid-increment *d*_0_ = (*d*_1_ + *d*_2_)/2 symmetrizes the problem so that *d*_l,2_ = *d*_0_ ± Δ*d*/2. Thus, we can analyze the estimation precision around the baseline distance *d*_0_ and decide whether Δ*d* can be discerned depending on the uncertainty of the distance estimate.

Let *σ*_*d*_ denote the one-dimensional standard deviation of any unbiased statistical estimator of the distance between the fluorophores. For such an estimator, the Cramer-Rao lower Bound (CRB) asserts *σ*_*d*_ ≥ *σ*_CRB_, where *σ*_CRB_ is obtained from the CRB. To state “resolvability” as the ability to safely distinguish two distances, we define the dimensionless ratio

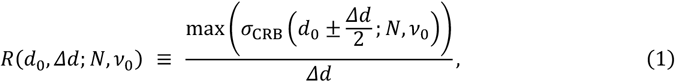

with *N* denoting the number of photons detected from each scatterer. A small change Δ*d* is defined to be resolvable if *R* < 1, which is tantamount to the maximum uncertainty of the estimate being smaller than the change in distance (Fig. 2A). *R* can also be understood as the maximum magnitude of the relative error of the estimate. For a fixed baseline distance (such as *d*_0_ = 0.05*λ* = 25 nm for *λ* = 500 nm) and a fixed relative change in distance Δ*d*/*d*_0_, the resolution scales with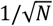 (Fig. 2B). Applying this criterion to the CRB of the distance of two molecules yields a simple inequality for the resolvability of a change in distance:

**Fig. 2.**
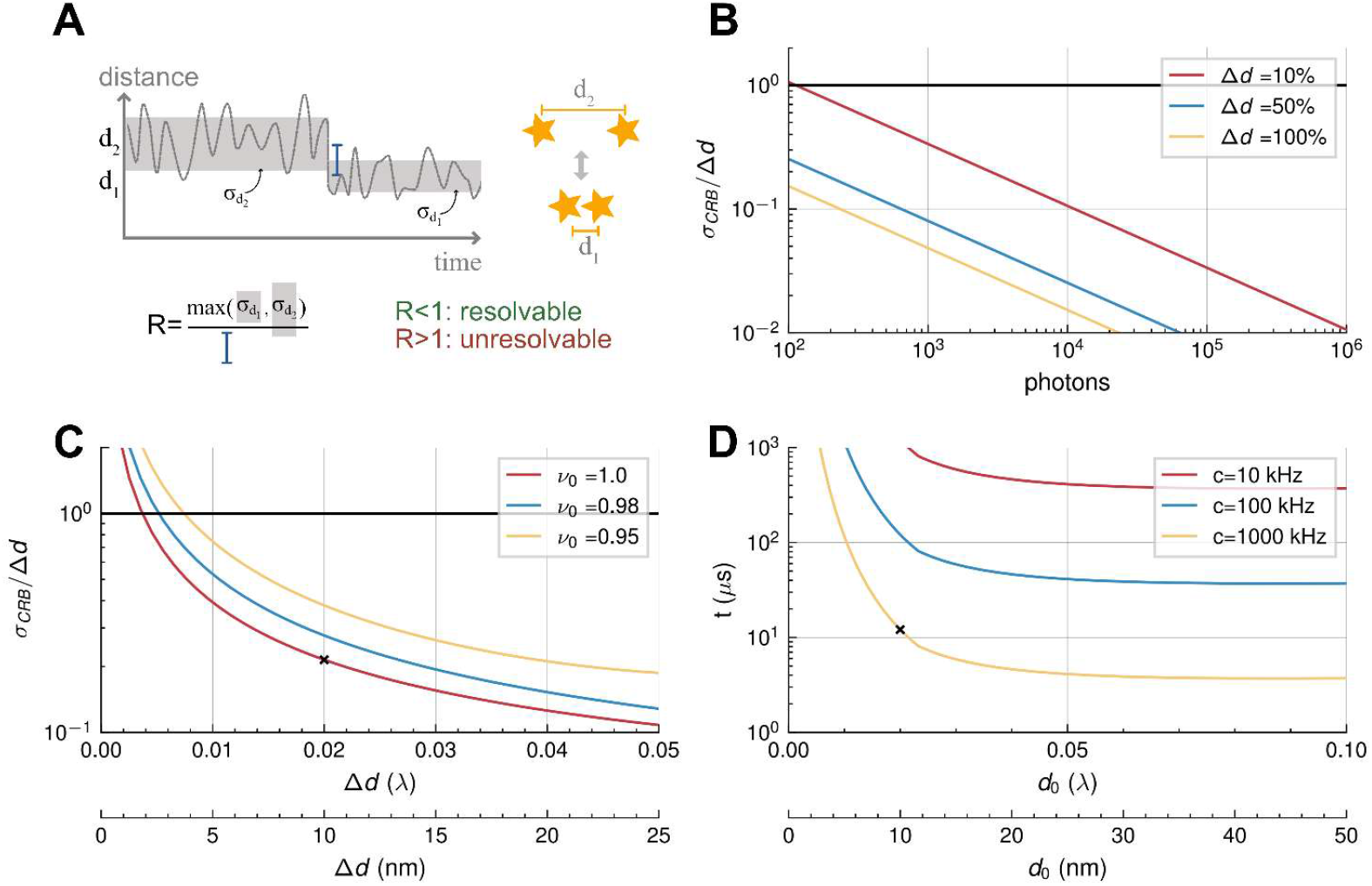
Theoretical limits of the spatio-temporal resolution when jointly tracking and resolving two identical incoherent point scatterers with an illumination minimum. (**A**) Schematic of two scatterers (stars) changing their distance from *d*_2_ to *d*_1_. The criterion R sets a lower bound on a change in distance *Δd* = |*d*_2_ − *d*_1_ | that can be distinguished from *d*_0_ = (*d*_1_ + *d*_2_)/2. (**B**) Relative precision of detecting *Δd* as a function of the number of detected photons. The baseline distance is *d*_0_ = 0.05*λ*, whereas *Δd* is stated as a percentage of *d*_0_. A minimum of zero intensity of the excitation beam is assumed, i.e., perfect contrast *ν*_0_ =1. (**C**) Relative precision of detecting *Δd* for three contrast values *ν*_0_ of the excitation beam and *N* = 100 collected photons per individual scatterer. To exemplify concrete distances, the wavelength in the sample medium is set to *λ* = 500 nm meaning that *d*_0_ = 25 nm. For *ν*_0_ = 1, a change Δ*d* = 10 nm can be measured with a relative error of 20 %. (**D**) Acquisition time required to resolve a 50 % change in distance depending on the baseline distance *d*_0_ for three different rates *c* of detected photons per single scatterer and *ν*_0_ = 0.99. At *c* = 1 MHz and *d*_0_ = 10 nm, a change *Δd* = 5 nm is identified in 12 *µ*s.

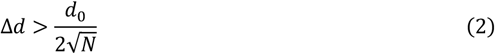

Let us now consider a baseline distance *d*_0_ between 0 and 5 % of *λ*, which in our numerical example would amount to 0 - 25 nm, and recall that the distance is encoded in the depth of the fluorescence signal minimum. If the minimum of the excitation beam is non-zero (imperfect contrast *ν*_0_ <1) the resulting fluorescence minimum is non-zero as well, making co-localization more uncertain due to the non-vanishing shot noise of the minimum of the signal. The minimally resolvable change in distance obviously depends on the initial contrast of the excitation intensity and the number of detected photons. For *N* = 100 photons per fluorophore and a baseline distance of *d*_0_ = 0.05*λ* = 25 nm, the smallest resolvable change in distance is ∼ 0.0075*λ* = 3.75 nm for an imperfect contrast *ν*_0_ = 0.98. For a perfect contrast *ν*_0_ = 1, the smallest resolvable distance change is ∼ 0.005*λ* = 2.5 nm (Fig. 2C).

Stating the resolution in terms of the number of collected photons allows us to obtain an expression for the temporal resolution with which a change in distance can be measured. Given a count rate *c* in units of photons per time interval, the number of collected photons during time *t* is *N* = *c* × *t*. The root of the equation *R*(*t*) = 1 is numerically found, yielding the time *t*_min_ needed to reduce the uncertainty of the distance estimate so that *R* ≤ 1. For any *t* ≥ *t*_min_, the threshold *R* = 1 is fulfilled, meaning that Δ*d* is resolvable:

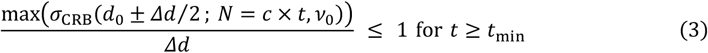

Depending on the fluorescence count rate of typically 10-1000 kHz, a 50 % change of a baseline distance *d*_0_ ≥ 0.02*λ* ≈ 10 nm is detectable within a time span of a fraction of a millisecond down to few microseconds (Fig. 2D).

### Quantifying the change in the ‘center-of-mass’ of two scatterers

While the relevance of establishing the position of the COM is highly evident, it is less evident how useful it is to quantify the precision with which the COM can be localized. This usefulness stems from the remarkable fact that, in our MINFLUX-based measurement, (sudden) changes in distance between the fluorophores lead to (sudden) changes in localization precision of the COM. Concretely, if *d* suddenly increases, the precision of the COM localization suddenly decreases, and vice versa.

An intuitive explanation for this relationship is that densely packed fluorophores ‘fit’ much better into the illumination minimum than fluorophores that are further apart from each other. Increasing their distance just increases their exposure to regions of higher illumination intensity, thus elevating the minimum of their joint fluorescence signal (Fig. 1D). Hence, the minimum becomes noisier and the COM localization less precise (Fig. 3A, B).

**Fig. 3.**
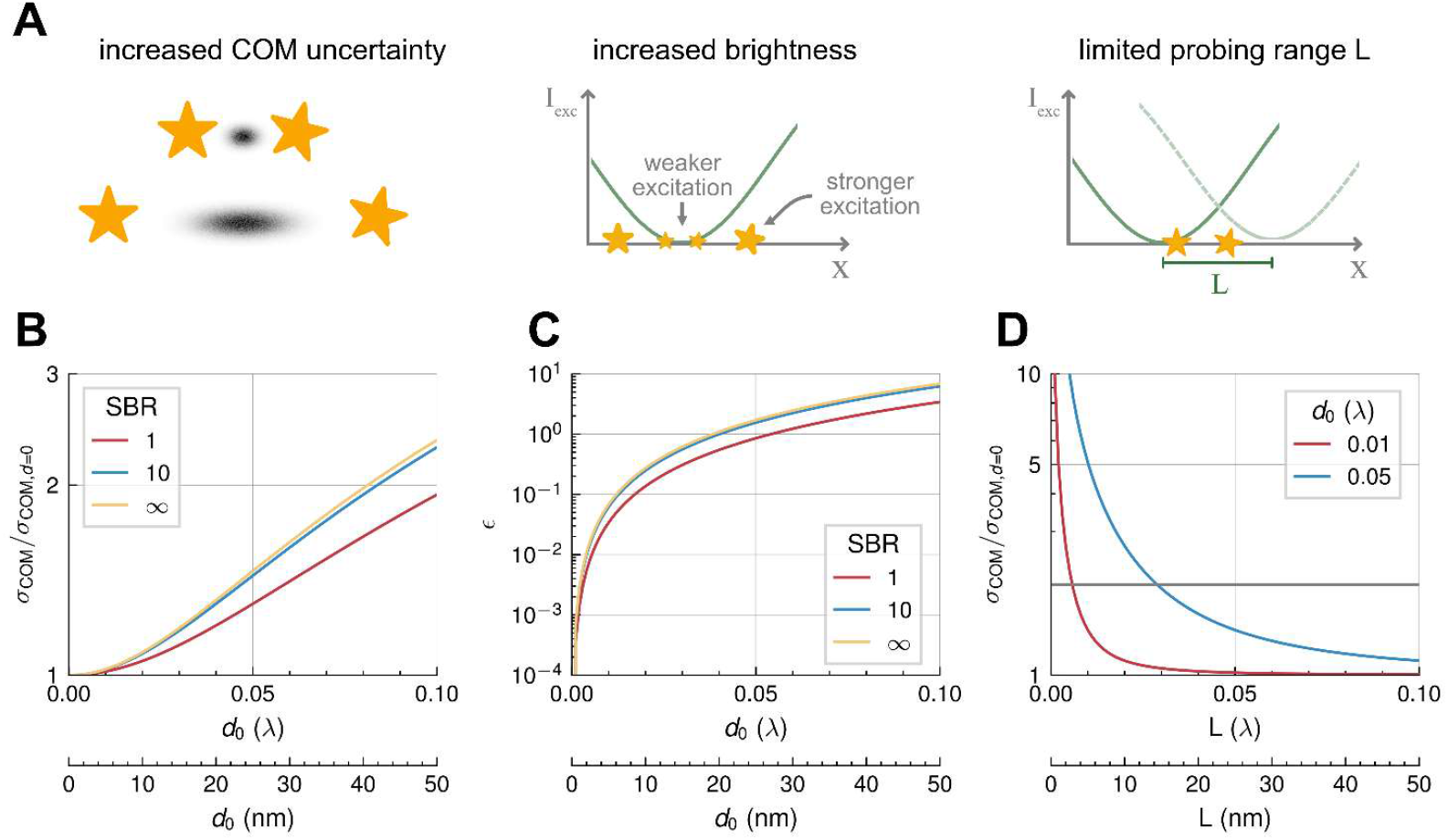
Theoretical limits on the center-of-mass (COM) localization precision of two incoherent point scatterers. (**A**) Sketches indicate that increasing the distance between the scatterers decreases the precision of their COM localization (left) and brightness (center), and also places a lower bound on the acceptable probing range L due to the inherent noise of the non-zero signal minimum. (**B**) The localization precision of the COM of the two scatterers deteriorates with their distance, compared to *d*_0_ = 0; the signal-to-background-ratio (SBR) is chosen as parameter. (**C**) Increasing *d*_0_ increases the relative brightness *ϵ* over the brightness at *d*_0_ = 0. (**D**) Given a baseline *d*_0_, the relative precision of the COM localization deteriorates with decreasing scanning range L, setting a lower bound on the applicable L. Avoiding precision decrease by >2-fold requires scanning ranges *L* > 6 nm and *L* > 20, for *d*_0_ = 5 nm and *d*_0_ > 25 nm, respectively (gray horizontal line).

To quantify this relationship, let *σ*_COM,*d*_ and *σ*_COM,0_ be the standard deviation of the COM estimate at a finite separation *d* and *d* = 0, respectively. Assuming the same photon statistics, we now define a signal-to-background ratio (SBR) as the ratio between the fluorescence signal at the minimum and any non-signal-related background. The factor showing the increase in localization uncertainty with increasing *d* is now given by (see SI Appendix, Methods)

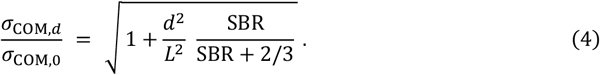

Additionally, we note that the brightness of the system changes with increasing distance *d*. The relative brightness at constant input power is found to be 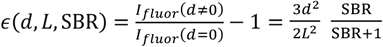, i.e. it grows with increasing separation *d* (see SI Appendix, Methods).

Assigning the correct specific positional states of the two-fluorophore system to an experimentally recorded fluorescence time trace yields the spatial evolution of the states, including their duration and the background. State changes can be identified through the changing depth of the minimum of the fluorescence signal, as well as through the changes in position and positional spread of the COM. The COM position is readily evaluated by fitting a Hidden-Markov-Model (HMM) with Gaussian emission probabilities to the COM time series. This approach can be further refined by taking into account the brightness of the two fluorophores via a Poissonian component of the HMM. In fact, it is expected that the fluorescence detection rate increases with increasing separation, as the fluorophores dive into regions of higher excitation intensity (see Fig. 3A, C).

Note that this behavior poses a challenge to our separation by MINFLUX, as unexpected changes in brightness of individual fluorophores may be confused with changes in distance. While total brightness variations can be calibrated from single-fluorophore data, individual brightness variations are not easily disentangled from temporal changes in distance. A potential remedy is to replace the point-like (confocal) detector used so far in our method with a camera, because by offering a spatially structured sensitivity in the detection plane, a multi-pixel detector can reveal brightness differences between the fluorophores.

Localization with a diffraction illumination minimum also overcomes the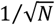 shot-noise scaling of the localization precision through iterative reduction of the probing range *L*. This well-known characteristic of MINFLUX single-fluorophore localization also applies to the simultaneous localization of two point-scatterers. For single fluorophores, the strategy of continuously reducing *L* to zero to improve precision is only constrained by an imperfect contrast *ν*_0_ < 1 of the illumination pattern and by a non-zero background, since both imperfections lead to a finite and hence noisy signal minimum.

For two equally bright emitters with finite distance *d*_0_ ≠ 0, however, the minimum of the joint fluorescence signal, i.e. at their COM, is always non-zero. This applies even for *ν*_0_ = 1 and zero background. Therefore, the localization precision of their COM is always poorer than that of two fluorophores with *d*_0_ = 0. Since identifying the non-zero minimum by scanning requires a certain signal from the slope of the minimum, the probing range *L* inevitably has a lower bound depending on *d*_0_. Reducing *L* for *d*_0_ ≠ 0 can easily increase the localization uncertainty of the COM by an order of magnitude over the localization precision obtained for fluorophores with *d*_0_ = 0 (Fig. 3A, D). Therefore, one typically imposes a limit on the ratio *Γ* = *σ*_COM,d_/*σ*_COM,0_, such that the COM localizations remain reasonably confined. This yields an explicit lower bound on the sampling width *L* via equation (4):

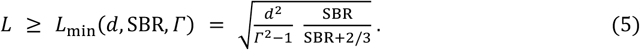

For any chosen *Γ* an optimal *L* has to be found that improves the precision of the distance estimate, but does not overly deteriorate the localization precision of the COM. Considering these limitations, our results still imply that it is possible to co-track two simultaneously emitting fluorophores on a millisecond or even microsecond timescale with a resolution sufficient to observe spatial dynamics on the order of a few percent of the employed wavelength.

### Resolving the positional states of a DNA nanostructure-based molecular switch

To mimic the conditions presented by a fluorophore-labeled protein or other biomolecule that changes its conformational state and hence the coordinates of the fluorophores bound to it, we took advantage of the nanometric precision and flexibility provided by the DNA-origami technique (8, 9). Concretely, we designed a DNA origami sheet with one of two fluorophores rigidly bound to the body of the origami immobilized on the cover slip, whereas the second fluorophore was attached to a ∼16 nm long double-stranded DNA lever arm. The lever was connected to the body of the origami via a flexible single-stranded linker, allowing it to move by Brownian diffusion. This way, the second fluorophore was able to explore the half-sphere above the cover slip. To prepare well-defined distances between the two fluorophores, the DNA origami sheet featured two binding sites capable of weak basepairing with the DNA lever, causing a transient arrest of the second fluorophore. The binding-sites were 23 nm apart, as well as 16.1 nm and 16.4 nm (one base-pair difference) away from the anchor point of the arm (Fig. 4A).

**Fig. 4.**
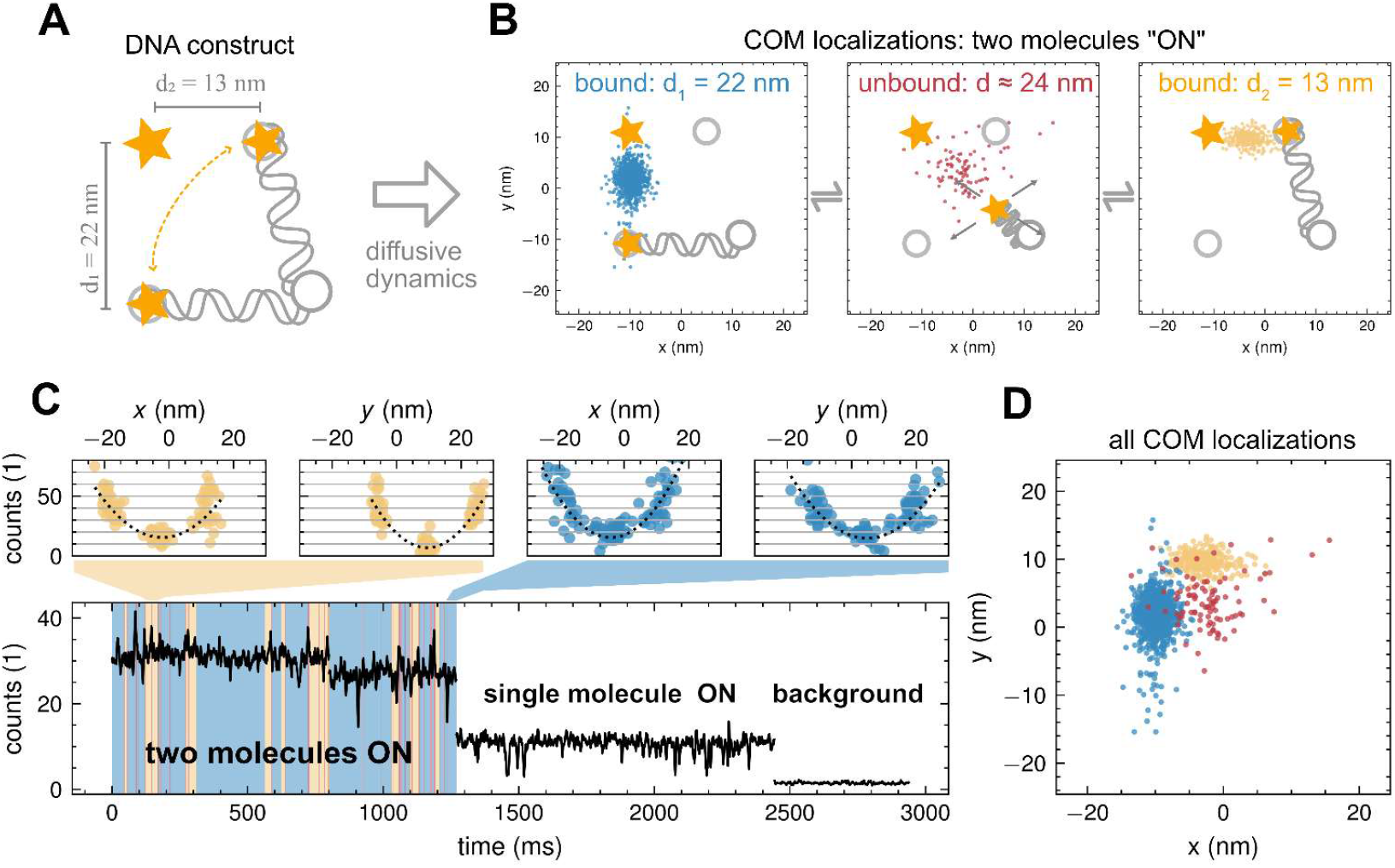
DNA-origami based construct (**A**) to measure quickly varying distances *d*_l,2_ between a fixed fluorophore (upper left star) and a movable fluorophore (lower star with indicated positional changes). A double-stranded piece of DNA with one fixed end (lower gray circle) and a fluorophore bound to its movable end entails two distinct distances between the fluorophores. Whereas the fixed fluorophore is at a specific position at the cover slip origami, the movable DNA end transiently binds at two different positions (upper gray circles). Driven by diffusion, the DNA strand randomly moves between the two bound states, yielding distances *d* = 13 nm and *d* = 22 nm between the fluorophores. (**B**) The localization routine identifies the center-of-mass (COM) of the two fluorophores. As the DNA strand moves, the identified COM positions assume different distributions (colored dots), allowing to infer the actual binding site of the movable strand, i.e., the positional ‘state’ of the construct, and assign this state to each individual localization based on a Hidden Markov Model (HMM). This reveals a short-lived unbound state discovered during the analysis. (**C**) Top: Probing the sample at three positions around the current COM reveals the typical parabolic dependency of the counts on the scan position. The position of the minimum informs the next choice of the center probing position, curvature and offset change depending on the distance and orientation of the underlying two fluorophores. Bottom: The averaged photon counts (time-track) over the course of measurement exhibits two distinct bleaching steps, proving an initial two-fluorophores stage, followed by a one-fluorophore and a background stage. The color-shading indicates the positional states of the construct, as inferred from the COM localizations. The single-molecule trace can be either due to the static or the mobile one, depending on which one bleaches first in the particular trace. The background is recorded for calibration purposes. (**D**) Scatter plot of the positions of the fluorescence minimum representing the COM of the two fluorophores over time. Three spatially distinct states are clearly visible, corresponding to the two bound and an unbound state (red). The segmentation is obtained by modelling the COM localizations with an HMM. In this specific sample, the unbound state is short-lived with a median dwell-time of 2 ms, while the bound states marked in blue and yellow exhibit dwell-times of 26.5 ms and 13 ms, respectively. The COM localizations are also depicted in the sketch in B for convenient comparison.

This macromolecular system consisting of a fixed and a quickly moving fluorophore offered three positional states: two states in which the moving fluorophore and hence both fluorophores were bound and a mobile state where the arm and hence one of the fluorophores could explore the available volume by diffusion (see Fig. 4B). As the two binding sites differed in distance and orientation with respect to the fixed fluorophore, this DNA construct mimicked three ‘conformations’ with different orientations in space and different inter-fluorophore distances. The states with both fluorophores bound implied either *d*_1_ = 22 nm or *d*_2_ = 13 nm.

Right after the arm became arrested at a binding site on the cover slip, the two-fluorophore system was expected to remain stationary for the duration of the binding event. The specific duration depended on the origami design, such as the available number of bases of the single-stranded binding segment at the sites and the arm, as well as on the salt concentration of the buffer, the GC content and the temperature.

Tracking the position of the two fluorophores and their COM was accomplished by placing the x- and y-oriented line-shaped focal intensity minima sequentially to three distinct positions of a spatial interval including the two-fluorophore systems as in the previously described MINFLUX tracking of single fluorophores. This positioning algorithm was reportedly more accurate than continuous line-scans in the x-and y-direction. They also allowed for relatively high update frequencies. Concretely, the fluorescence photon counts were probed for a total of six points, by positioning the *y* -oriented line-shaped minimum along the *x*-axis at -15 nm, 0 and 15 nm relative to the last COM estimate. Instantly thereafter, the *x*-oriented line-shaped minimum was equally placed along the *y*-axis.

Determining the fluorescence minimum by an independent parabolic fit along each axis led to a new estimate of the COM (Fig. 4C). To capture the characteristic transition times of the DNA construct, the acquisition time per probing point was chosen so that the total time per COM estimate was 0.63 ms or 0.98 ms. The sum of all six recorded photon counts formed a time trace with discrete bleaching steps, revealing the number of emitting fluorophores (Fig. 4C). To achieve sufficient precision per time step without accelerating fluorophore bleaching, the laser power was adjusted such that the detected photons per fluorophore ranged between 15 and 50 per scanning position.

In order to co-track the two molecules and ultimately retrieve the conformational states of the DNA construct, we first identified the COM of the two fluorophores. The COM occupied different positions (Fig. 4B), depending on the position of the arm. Tracking the COM eventually led to bleaching of one of the fluorophores, leaving most measurements with a pronounced single-emitter phase produced by the static or the movable emitter (Fig. 4C). After bleaching of the remaining molecule, the local background was recorded for calibration purposes.

Since the conformational changes of our DNA construct were pronounced (Δ*d*/*d*_0_ ≈ 0.5), we were able to segment the fluorescence trace just by observing the x- and y-coordinates of the identified COM (Fig. 4C, D). Being identical with the signal minimum of the two fluorophores, the COM localizations clearly featured distinct positional jumps and stationary phases. Applying an HMM, classified the COM localization as spatially well-separable clusters, revealing the conformational changes of the DNA construct during both the two- and the single-fluorophore phase (see Fig. 4D). The spatial arrangement of these clusters compares well with the expected positions of the COM of the three states of the construct (Fig. 4B, D).

The thermally driven movement of the DNA arm was interrupted only by binding to the two sites on the origami sheet. Hence, we expected the arm to assume only two distinct positional states. We also expected the fluorophore to diffuse so quickly that a position between two binding events is not measurable. Surprisingly, we were able to also localize the unbound state of the arm as an average COM position. The duration of this unbound state was on a millisecond time scale. Interestingly, the unbound COM position was found relatively far away from the fixed fluorophore, shifted away from the connecting line between the two binding sites towards the anchor point of the arm, consistent with electrostatic repulsion between the DNA arm and the DNA origami sheet. Therefore, we concluded that the arm stood out of the x,y-plane during diffusion. Altogether, the observation of the COM evolution already provided a basic understanding of the dynamics of the DNA construct, yet without providing the desired fluorophore distances.

In order to investigate this unexpected behavior and to verify the underlying macromolecular structure of our DNA construct, we analyzed the single-molecule part of the recorded traces. By selecting traces with at least one bleaching step, we identified traces where the mobile fluorophore attached to the arm remained active and was tracked by MINFLUX. Next, we determined the single-molecule positions in these traces from the position of the intensity minimum (see Fig. 4C). Averaging over many single-fluorophore traces of many constructs provided clusters of single-fluorophore localizations with sufficiently precise positional information, and thus measured reference distances between the two fluorophores.

Concretely, we expected three distinct positions of the movable fluorophore, namely two for the DNA arm bound and one for arm unbound (Fig. 5A). We also expected the fluorophore position to be more confined and longer-lasting in the two bound cases. Since the binding sites were arranged at equal distance from the arm’s anchor point, an unbiased movement of the arm was expected to produce similar distances between the averaged localizations of the fluorophore on the unbound arm to its averaged localization at the two binding sites.

**Fig. 5.**
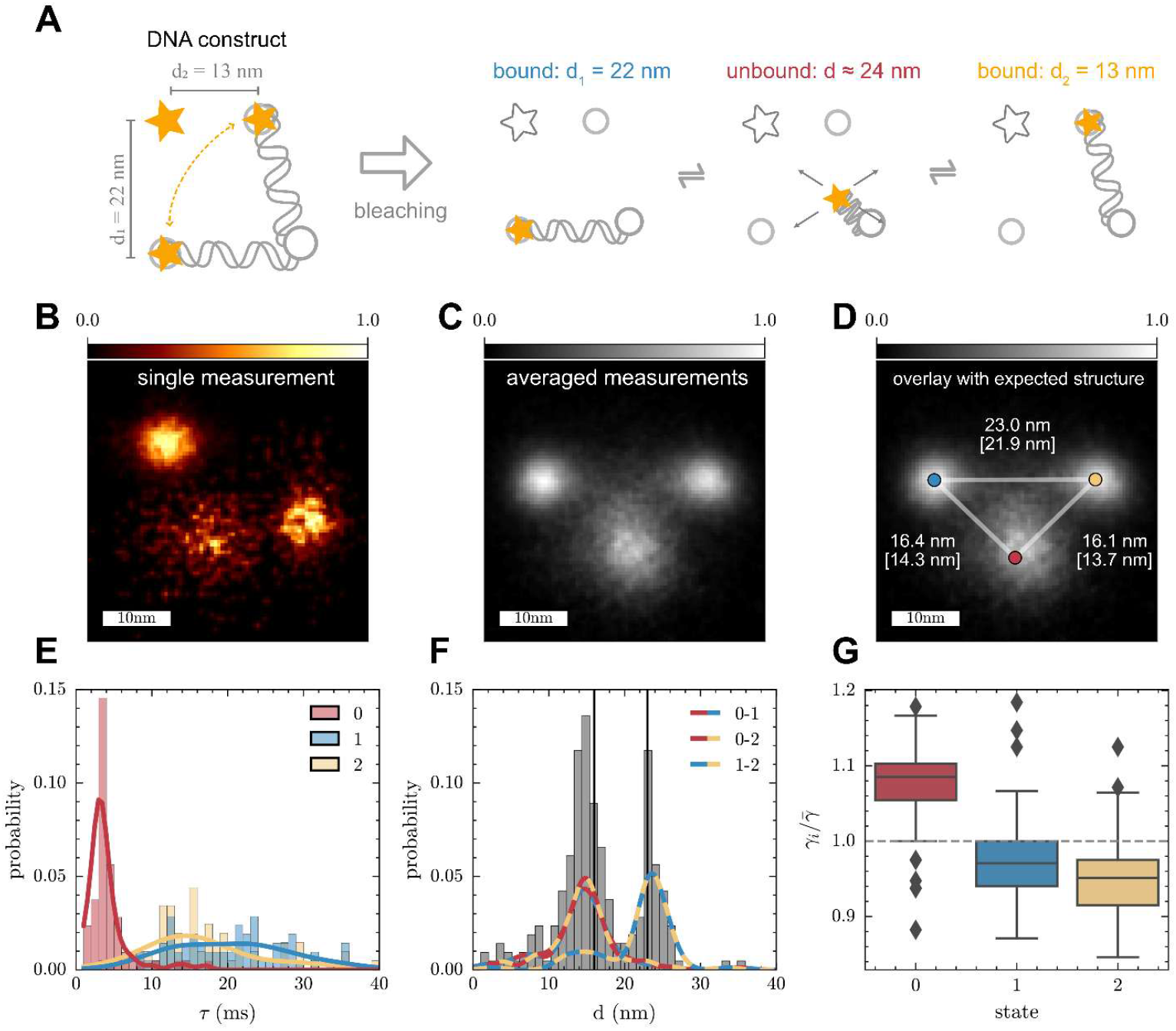
Tracks of the single fluorophore on the movable DNA arm confirm the design of the DNA construct with nanometric precision. (**A**) The fluorophore (filled star) has two bound (blue and yellow box) and an unbound position (red box) that are associated with the three ‘conformational states’ of the origami. They feature a distance of 16.4 nm and 16.1 nm between the anchor point of the arm and the binding sites, as well as a 23 nm distance between the binding sites. (**B**) Heatmap of the single fluorophore localizations from a single exemplary origami construct (60 photons/localization on average) shows three distinct clusters of localization. Given unbiased diffusion of the DNA strand, the movable fluorophore covers a hemisphere with the anchor point at its center (**C**) Average over the localization data of 70 identically designed origami constructs. For improved visualization, the localizations are normalized to unity peak density per state. (**D**) A least-squares based overlay of the expected positional arrangement of the anchor point (red) and the two binding sites with the averaged localizations and expected distances. In brackets: mean pairwise distances ± standard errors between the clusters are: 14.3±0.6 nm (for state 0 to 1), 13.7±0.5 nm (for state 0 to 2), and 21.9±0.6 nm (for state 1 to 2). (**E**) Dwell-times for the three spatially distinct states of 70 traces of the movable fluorophore. Each count in the histogram represents the median dwell time of a state in one of the traces; color code as in panel a. (**F**) Pairwise distances between the localization clusters of 70 traces. Distances are displayed in an unspecific histogram (gray) with state-specific kernel density estimates (dashed lines). The distance between the binding sites peaks at 23 nm, as expected (right black vertical line). The distances from the anchor point to either of the binding sites are equal but slightly shifted to smaller values than the expected 16 nm (left vertical line). (**G**) Relative brightness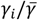 (ratio of the brightness of molecule at position 0, 1 or 2 and the average brightness among all positions) at the binding sites shows that the arm in the unbound state appears brighter, as expected from a fluorophore that explores a larger fraction of the intensity profile around the minimal intensity point. The other two binding sites exhibit a relative difference in median brightness of about 2%. The colored boxes signify the quartiles of the distributions with its median indicated by the horizontal line, the whiskers extend to show the rest of the data up to a 1.5 inter quartile range (IQR); rhombes show outliers beyond the 1.5 IQR.

To extract characteristic dwell-times and separate the localization clusters, we segmented the single-fluorophore traces with an HMM and assigned specific positions to each localization (see Methods). Since the mobile fluorophore was expected to assume three positions, we selected the traces displaying three distinct positions and flagged the position with the shortest dwell-time as the ‘unbound’ one. Next, we translated and rotated the localizations such that the short-lived position resided near the origin and the bound positions to the left (position 1) and the right (position 2) of the y-axis. This state-assignment was unambiguous, since the DNA arm had to lie face-up on the coverslip in order to be mobile, leaving translation and rotation as the only possible spatial transformations. Thus, we created a unique position mapping relative to the short-lived position, without having to consider the distances between the localization clusters at this point.

To combine the information from all 70 traces with a mobile single fluorophore into a common reference frame, we implemented a rigid averaging strategy, akin to the sub-tomogram averaging routines in electron microscopy. Each trace (Fig. 5B) was treated as resulting from a fixed configuration of three fluorophore positions, namely two for the arm bound and one for the arm unbound. The centroids of these positions were aligned by optimizing a rotation and translation with respect to a global template. This procedure did not assume a prior model for the positions; the consensus positions iteratively emerged from the data. Starting from a robust initialization based on orienting the two long-lived positions along the x-axis with the short-lived position below, the algorithm alternated between two steps: (i) fitting each trace to the current global template using a weighted Procrustes transformation, and (ii) updating the template as the weighted average of all aligned traces. The weights were chosen such that long-lived and well-localized positions contributed more strongly. After convergence, all traces were consistently registered in a common coordinate system, yielding a denoised representation of the average three positions (Fig. 5C, D).

The alignment procedure also allowed us to obtain ensemble-level statistics of the dwell times, pair-wise distances, and the relative brightness of the positions assumed by the movable fluorophore (see Fig. 5E-G). The short-lived unbound position featured a median duration of a few milliseconds, whereas its bound counterparts lasted for 10-30 ms (Fig. 5E). The pair-wise distances between the positions revealed two pronounced peaks: one at 14 nm and the other at 22 nm (Fig. 5F). The precise pairwise distances of the position medians after the particle averaging actually were established as follows 0-1: 14.28 nm; 0-2: 13.74; 1-2: 21.94 nm. The distances between the unbound to the bound positions (0-1 and 0-2) were equal, as expected. However, they were slightly smaller than the expected distances from the binding-sites to the anchor point of the arm; compare peak of kernel-density-estimates to vertical line at 16 nm, implying that the arm was slightly biased, “leaning forward” towards the binding sites. In other words, the arm neither laid flat on the origami sheet between the two binding sites nor did it stand perfectly upright, since the localizations were not distributed around the anchor point. The arm rather explored a conical volume, slightly leaning towards the two binding sites. Last, the separation of the binding sites (1-2) with 22 nm was close to the expected distance of 23 nm.

We also observed a slight imbalance in the relative brightness of the fluorophore for the different positions (Fig. 5G). By normalizing its median brightness in either of the three positions to the average median brightness of all positions, we noticed that the fluorophore was generally brighter when unbound (compare (10)). In the two bound positions, the fluorophore appeared to be similarly bright, with a difference of only a few percent. Altogether, the single-fluorophore traces fully validated the expected design of the DNA construct.

Therefore, we moved on to the decisive experiment, namely the simultaneous tracking and resolution of the two fluorophores. Note that the DNA origami construct has a size of about 3% of the employed excitation wavelength, which is tiny compared to the extent of the focal excitation pattern produced by diffraction (see Fig. 6A). The separation of the two emitting fluorophores was 13 nm and 22 nm for the bound states and about 24 nm for the unbound state (see Fig. 6B). With regards to the theoretical treatment, this value amounted to a baseline distance of *d*_0_ = 17.5 nm ≈ 0.03*λ*. The change in distance was *Δd* = 9 nm = 0.018*λ* ≈ 0.5*d*_0_. Using the fluorophore Atto647N, we achieved count rates on the order of 100 kHz and slightly higher. For these values, the numerically obtained lower bound on the temporal resolution was < 2 ms for baseline distances down to 0.02*λ*, and below 10 ms for distances down to 0.01*λ*. This finding suggested that temporal bins on the order of a few milliseconds should enable us to resolve both the two long-lived bound states and the short-lived unbound state.

**Fig. 6.**
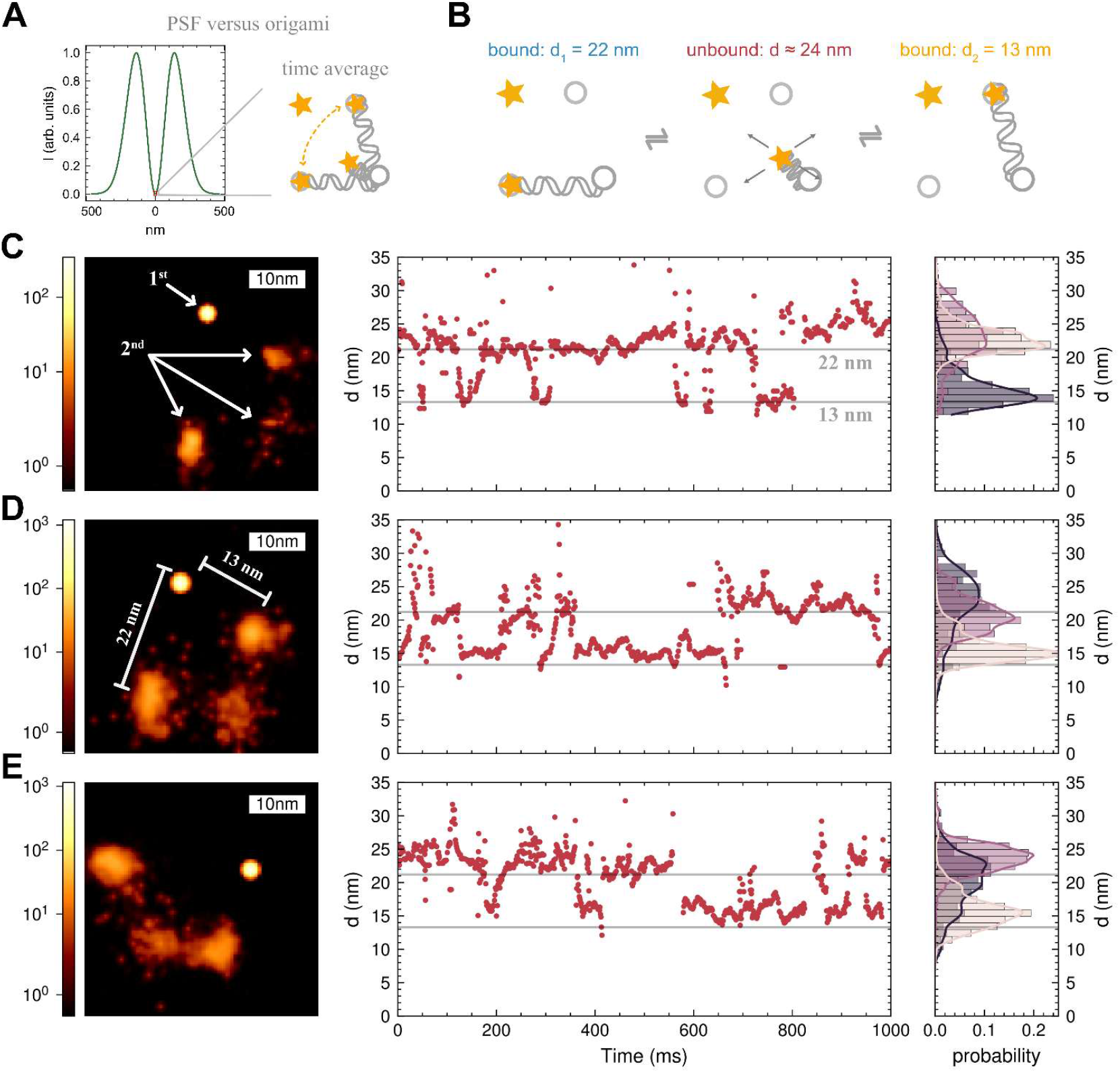
Simultaneous tracking and resolution of two fluorophores reveals the three conformational states of the DNA construct. (**A**) Illumination intensity profile drawn to scale to illustrate the position of the two identical fluorophores (stars) with respect to the minimum; their distance *d*_0_ ≈19 nm corresponds to 3 % of the employed wavelength *λ* = 640 nm. Note that MINFLUX utilizes the very minimum of the illumination beam. (**B**) The three conformational states of the two-emitter system: i) bound with 13 nm separation of the two fluorophores, ii) unbound state with ∼24 nm and iii) bound with 22 nm separation. (**C**) Left: heatmap of the number of simultaneous localizations of the static (1^st^) and the movable (2^nd^) fluorophore averaged over time, concomitantly demonstrating the nanometric resolution under constant joint emission. The static fluorophore’s position is estimated with high precision. Three distinct localization clusters mark the position of the movable (2^nd^) fluorophore. The motility of the 2^nd^ fluorophore yields more spread-out clusters, reducing the precision. Center: the change in distance between the fixed and movable fluorophore over time reveals two plateaus corresponding to the distances in the two bound states (horizontal gray lines). The trace directly reveals the temporal evolution of the conformational state of the DNA construct. Right: distance histogram shows two narrow peaks and a broader peak, corresponding to the bound states and the unbound state, respectively. (**D-E)** Two further measurements on different origami constructs as in C.

We localized the two fluorophores simultaneously using a Maximum Likelihood Estimate (MLE) of their positions. Accumulating these localizations over time yielded distinct clusters of localizations for each of the molecules (see Fig. 6C-E (heatmaps)). If available, the position of the fixed (1^st^) fluorophore was soft-constrained with a prior obtained from the single-molecule phase of the trace (see Methods). We note that this procedure allowed to improve the quality of the estimate of the movable (2^nd^) molecule, which jumped between the two binding sites and the unbound state. Yet, soft-constraining was not strictly necessary.

Plotting the inter-fluorophore separation over time revealed plateaus at 13 nm and 22 nm, as expected (Fig. 6C-E (center)). Here, a sliding-window of 15 ms and a stride of 2 ms was used to obtain the distance estimates. Sorted by the states obtained from the HMM, the obtained separation showed pronounced peaks around the expected distances (Fig. 6C-E, histograms). Note that Fig. 6C-E displays examples where the mobile fluorophore was never observed alone. The temporal evolution of the accumulated localizations and distances show that the two fluorophores were continuously tracked (see Movie S1, SI Appendix). Moreover, our approach consistently and correctly revealed their distances and orientation in the focal plane.

## Discussion

We showed that two incoherent point scatterers located within rapidly changing nanometric distances can be tracked and resolved without ever observing one of them individually. The diffraction resolution barrier was overcome by the MINFLUX approach, i.e., rapid spatial probing (scanning) across both scatterers with a diffraction illumination minimum and analyzing the associated spatial modulation of their joint signal. Thus, given the position of the minimum, the resolution problem was reduced to the estimation of the positions and brightness of the scatterers. Our approach radically departs from current fluorescence super-resolution methods, which, by requiring every fluorophore to be sequentially turned ON and OFF, are unable to provide continual fluorophore tracking with nanometric resolution.

The dynamic DNA origami construct used in our experiments was shown to undergo conformational transitions and randomly occupy three distinct conformational states: two in which the mobile part of the construct was bound and one in which it was unbound. The three states were recovered from the joint signal of two fluorophores with identical, non-changing spectral properties. In consistence with the theoretical limits derived, our experiments demonstrate that resolving the spatial dynamics of identical fluorophores at nanometer distances is feasible with the photon detection rates available in practical fluorescence microscopy.

Our findings raise the question of how our separation and distance measurement compares to conceptually straightforward methods that use dissimilar fluorophores for separation. In fact, excitation and/or emission disparities are widely used for measuring inter-fluorophore distances, either through individual localization or through fluorescence resonant energy transfer (FRET).

As a dipole-dipole transition scaling with *d*^−6^, FRET is highly responsive to changes in *d* (11, 12). However, its application is effectively restricted to the *d* = 2 − 8 nm range (13). The sensitive dependence of FRET on the orientation of the two dipoles makes accurate distance measurements with FRET difficult, rendering FRET more suitable as a binary indicator of close molecular proximities. Moreover, FRET does not yield the actual three-dimensional (3D) coordinates of each fluorophore, but just the distance *d* as a scalar. Changes in 3D-coordinates that maintain *d* as a value are not detectable, though we note that a combination of FRET and an approach called pMINFLUX (14) has enabled separation by fluorescence lifetime and precise tracking in and outside the FRET range (15).

While the particular MINFLUX setup used in our study provided only measurements in the x-y-plane and planar projections, more elaborate setups deliver 3D localizations (16). Therefore, dedicated MINFLUX systems should yield vectorial 3D-distances from a few nanometers up to ∼100 nm. Larger distances are not challenged by diffraction. Also, the precision of our distance estimate scales linearly with *d*, making it more robust than FRET.

As a dipole-dipole interaction, FRET is difficult to calibrate, whereas the calibration of the excitation contrast *ν*_0_ used here is based on a simple least-squares-fit on single-fluorophore traces, which is a by-product of the recorded data. Last but not least, FRET cannot be readily scaled up to measurement schemes using more than two interacting fluorophores (17).

In fact, this problem holds for any multicolor single-fluorophore approach. Using multiple different fluorophores at multiple binding sites demands a highly selective and hence more complex biochemical labeling protocol. The need for perfect labeling stoichiometry, meaning that every macromolecule has to be labeled on every relevant site with a fluorophore of different color, massively complicates the labeling. Failure to meet specificity and completeness poses major challenges for the interpretation of multicolor data.

Even if a number of spectrally distinct labels are attached to a biomolecule with sufficient coverage, the broad and strongly overlapping excitation and emission spectra of organic fluorophores entail multiple cross-talk both in the excitation and in the detection process, challenging their separation in a multicolor detection setup. In fact, each emission ‘color’ in the detection path requires some tens of nanometers in spectral width in order to be separable with confidence.

The difficulty with multiple colors is exacerbated by the fact that, at the sub-10 nm scale, chromatic aberrations in the microscope’s optical train are difficult to gauge. Thus, chromatic aberrations limit the accuracy of localization of two or more fluorophores of different excitation or emission wavelength, challenging the measurement of absolute distances between individual fluorophores (18–21). By requiring only identical fluorophores and thus just a single-color excitation and detection channel, our MINFLUX separation is entirely monochromatic and hence by definition devoid of chromatic problems, which is an invaluable advantage in the sub-10 nm regime. In this context it is also important to realize that our method is not compromised by potential FRET between identical fluorophores, i.e., symmetrical homo-FRET, because in their joint signal the actually emitting fluorophore is irrelevant.

Nonetheless, MINFLUX critically depends on the depth of the intensity minimum of the illumination light. Making the minimum ‘deeper’ improves the separation capability, the localization precision and thus the detection of conformational states. An interesting strength of our method is that it benefits from having multiple emitters in close proximity, due to the fact that the signal of the multiple emitters ‘stack up’, making the minimum of their joint signal more readily distinguishable from background. This should be contrasted to the previously discussed multicolor conventional localization which is practically limited to 2-4 markers, due to the noise and background challenges from spectral cross-talk, making multicolor separation at small distances mathematically unreliable.

Although our approach also applies to longer distances and offers comprehensive positional information, its precision and temporal resolution does not yet match those obtained by FRET under optimal conditions. In the future, substantial improvements should also be obtained by applying more elaborate MINFLUX scanning patterns. There is no fundamental reason why our method could not be scaled down to distances much shorter than those reported in this study, once background and the contrast of the illumination minimum are improved. Like in the early development of 4Pi-microscopy (22), applying a near-infrared wavelength for two-photon rather than single-photon excitation is another viable option for drastically deepening the excitation minimum due to the quadratic dependence of the fluorescence on the excitation intensity (23). By the same token, the comparatively long near-infrared excitation wavelength facilitates the separation of visible fluorescence light and reduces background.

Nonetheless, two critical provisions of our method still require full attention. First, as it stands, separation by MINFLUX is only applicable to a known and fixed number of emitters. If the number of emitters is unknown, it needs to be determined, e.g., through analyzing the total brightness of the emitters, or counting the bleaching steps at the end of the measurement. A less straightforward but workable solution is to extract the number of emitters from the photon emission statistics (24). Spontaneous blinking, misclassified photobleaching, or labeling heterogeneity can lead to misinterpretation of the positional states and fluorophore distances. Second, separation by MINFLUX requires knowledge of the relative brightness of the emitters. Here, we have assumed the emitters to be equally bright. Fluctuations in brightness arising from excursions of the fluorophore to long-lived dark states or from interactions with the molecular environment may affect this assumption on certain time-windows. Since distance estimates are sensitive to variations in brightness, modelling the brightness as a latent variable (e.g., in a joint position-brightness HMM), may turn out to be necessary in more complex settings. Note that such brightness variations also affect established multicolor separation, including FRET.

We also remark that MINFLUX separation has so far been investigated for resolving incoherent, inelastic scatterers such as fluorophores only. Extending the method to elastic scatterers requires the consideration of both the scattered wave field and phase. Yet this expansion should be straightforward, since the phase differences between individual scatterers at nanometric distances are rather small.

In summary, we have shown that rapid scanning with an illumination beam featuring a zero-intensity node at a well-defined position is able to separate individual identical point scatterers, particularly fluorescence molecules, undergoing nanometric distance changes within milliseconds and less. This novel sub-diffraction resolution principle opens up an arguably unexpected route to exploring the inner workings of biomolecules, especially proteins, under native conditions. Once developed into a widely applicable method, this principle should allow the direct visualization of 3D-conformational changes of individual macromolecules in living cells.

## Materials and Methods

Details of the sample preparation, data acquisition and analysis can be found in *SI Appendix*.

## Data Availability

All study data are included in the article and/or supporting information.

## Author Contributions

TAH and SWH designed the research. TAH and OLS performed the research. TK and KG contributed the DNA construct. TAH analyzed the data with input from OLS and SWH. TAH and SWH wrote the paper.

