## Supplementary material for "Nonstop nanometric resolution of randomly moving point scatterers with focused light": SI Appendix

### Supporting Information Text

#### Materials and Methods

##### MINFLUX measurements

The MINFLUX tracking routine requires an initial COM estimate $x_{\mathrm{COM}}[0]$ of the fluorophores that have been attached to the DNA origami. This initial estimate is obtained by fitting a 2D-Gaussian to the area around the locally brightest spots in a Gaussian kernel smoothed confocal scan. The field of view of the confocal scans was 10 x 10 µm², with a pixel size of 50 x 50 nm², acquired with a dwell time per pixel of 100 µs and a 1-5 µW laser intensity.

The MINFLUX tracking routine is implemented by probing three fluorescence photon counts $[n_-, n_0, n_+]$ at positions $[-L/2, 0, +L/2]$ relative to $x_{\mathrm{COM}}[t]$ along the $x$ and $y$ direction. The new COM estimate is then calculated independently for each direction according to

$$x_{\mathrm{COM}}[t + \mathrm{d}t] = x_{\mathrm{COM}}[t] + \frac{L}{4}\frac{n_- - n_+}{n_+ + n_- - 2n_0},$$

which is derived from a parabolic fit to the three intensity values in a single direction. The position is not updated if fewer than five photons are collected, or if $2(n_+ + n_- - 2n_0) < |n_- - n_+|$, which limits the allowed absolute COM corrections to $L/2$ and excludes fits with negative curvature.

To account for the greater uncertainty of the initial COM estimates, a zoom-in approach was chosen. This started with an initial L of 200 nm, which was then reduced to 30 nm in a pre-programmed manner over the first ten localizations (3x $L$=200nm, 4x $L$=100nm, 3x $L$=60nm). The remaining localizations were acquired with an $L$ of 30 nm. The time per $xy$ localization $\mathrm{d}t$ was set to 0.63 ms or 0.98 ms to capture the typical transition times of the DNA origamis arm. The laser intensity was adjusted so that the sum of all photons emitted by a single fluorophore per localization was between 15 and 50.

##### Sample preparation

*Design of DNA origami structures*

The DNA origami structure was based on earlier designs by Kopperger et al. (1) and Dreher et al. (2) and was adapted using cadnano (version 2.4) (3). The structure uses the p8064 scaffold and was folded into the desired square-shaped two-layer sheet using 188 oligonucleotide staple strands. Staples at the outer edges were omitted to reduce stacking interactions between separate DNA origami structures (2). A complete list of staple sequences is provided in Table S1, and the cadnano design is shown in Figure S1.

On the top side of the two-layer sheet, two staples were extended to create binding sites for transient hybridization (binding site1/2; two thymidine spacers followed by the 7-nt sequence 5′-AGTCCTA-3′). These two sites were positioned to enclose a 90° angle with respect to a third extended staple carrying four thymidine spacers and a covalently linked Atto647N fluorophore (static fluorophore). A fourth staple, located at approximately equal distance from the two transient binding sites, was extended with three thymidine spacers followed by a 15-nt sequence forming the base of a flexible hinge (hinge base). Two additional strands completed the hinge: one carrying an internal Atto647N fluorophore (mobile fluorophore) and one containing a single-stranded sequence at the hinge terminus that is reverse-complementary to the transient binding-site sequence (hinge binding site). All modified strand sequences are listed in Table S2.

*DNA origami assembly*

DNA origami structures were assembled using 10 nM p8064 scaffold strand (tilibit nanosystems, Munich, Germany) in 10 mM Tris and 1 mM EDTA (TE buffer). Unmodified staple and hinge strands (Integrated DNA Technologies, Coralville, IA, USA; standard desalting) and fluorophore-labeled strands (biomers.net GmbH, Ulm, Germany; HPLC purified) were dissolved in TE buffer and added at a fivefold molar excess relative to the scaffold. The folding mixture contained 1× Tris–acetate–EDTA (TAE; Sigma-Aldrich) and 20 mM $MgCl_2$ (Sigma-Aldrich). The origami structures were folded by thermal annealing in a thermocycler (Bio-Rad C1000 Touch), cooling from 70 °C to 20 °C over 12 h, followed by a 3 h hold at 40 °C and subsequent cooling to RT.

##### *DNA origami purification*

Folded structures were purified from excess staples by spin filtration using an Amicon Ultra centrifugal filter (100 kDa MWCO; Merck Millipore, UFC5100BK) in a Fresco 17 microcentrifuge (Thermo Scientific, 75002420) at 4°C. Filters were pre-rinsed by centrifugation with 500 µL of 1× TAE and 5 mM $MgCl_2$ for 5 min at 13,000 × g. Samples were loaded in two rounds by combining 25 µL DNA origami with 475 µL of 1× TAE and 5 mM $MgCl_2$ and centrifuging as above. DNA origami was recovered by inverting the filter into a fresh microtube and centrifuging for 2 min at 1,000 × g. The $MgCl_2$ concentration was then adjusted to 20 mM. Samples were stored for up to 5 days at 4 °C until imaging.

##### *AFM imaging and analysis*

To confirm successful folding, purified DNA origami samples were imaged by high-speed atomic force microscopy (AFM). Two microliters of purified sample were diluted in 48 µL imaging buffer (1× TAE, 20 mM $MgCl_2$), pipetted onto freshly cleaved mica (Sigma-Aldrich, AFM-71856-02), and incubated for 1 min. Subsequently, 1 mL imaging buffer was added to the imaging chamber. Imaging was performed on a JPK Nanowizard Ultra Speed AFM (Bruker) in AC mode using a FastScan-D cantilever (resonance frequency in water: 110 kHz; spring constant: 0.25 N/m; Bruker).

Images were processed using Gwyddion (version 2.63, (4)) by applying standard operations such as flattening, removal of faulty lines and artifacts, and the manual adjustment of color scales.

##### *Cleaning process of the cover slips*

The coverslips (170 µm, No. 1.5H, 18 x 18 mm², Paul Marienfeld GmbH & Co. KG) were cleaned by incubating them in acetone, followed by isopropanol and then double-distilled water, for 5 minutes each. After removing them from the double-distilled water bath the coverslips were dried using a nitrogen flow. No more than five days before a measurement, the coverslips were plasma cleaned using oxygen for 5 minutes at 200 W.

##### *Construction of flow chamber*

A flow channel was made to allow the exchange of reagents and buffers during sample preparation. The flow channel was prepared by attaching two strips of scotch tape (Scotch double-sided tape, 3M) to a microscopy slide at a distance of about 5 mm. The channel was then closed by placing a previously cleaned coverslip on top. Finally, any excess tape was removed using a scalpel.

##### *Preparation of the DNA origami sample*

To prepare a sample for imaging, the flow chamber was first washed with TAE buffer and the concentration of $MgCl_2$ chosen for the measurement. A solution of the DNA origami was then flushed in and incubated for 5 min. The dilution of the DNA origami solution was chosen so that the final surface density of origami was about 0.1 $\mu m^{-2}$. After the incubation period the unattached origami were flushed out of the channel by adding the imaging buffer. The imaging buffer was based on TAE buffer with the corresponding concentration MgCl2, 10% glucose (D(+)-glucose monohydrate, cat. no. 6780.1, Carl Roth GmbH + Co. KG), 0.1 mM methyl viologen (cat. no. 856177, Sigma-Aldrich Chemie GmbH), 0.1 mM Trolox (cat. no. 23881, Sigma-Aldrich Chemie GmbH) and adding 1 µL of pyranose oxidase enzyme system per 100 µL of buffer. The pyranose oxidase enzyme system was prepared by solving 10 mg of pyranose oxidase (cat. no. P4234, Sigma-Aldrich Chemie GmbH), 170 µL PBS, 80 µL catalase (cat. no. 190311, MP Biomedicals). In the end the slide was sealed with either epoxy glue (UHU 2x Plus Sofortfest, UHU GmbH & Co. KG) or picodent twinsil speed 22 (Picodent Dental-Produktions- und Vertriebs GmbH).

#### Data analysis and localization

The analysis segments, filters and processes localization traces to obtain the temporal state assignments of the DNA structure, system-related quantities like PSF parameters (contrast and brightness of illumination) and finally emitter positions (co-localizations). In the following, we give a conceptual summary of our analysis routine.

In a first segmentation step, a series of position-count tuples is first standardized and partitioned into quasi-stationary blocks using a change-point detection algorithm (5). Each block represents a segment in which the emission statistics are approximately constant. The parameters of the change-point detection are empirically chosen such as to yield break points at the bleaching steps of a trace or significant changes in brightness.

Second, we filter the obtained segments and initiate an iterative labelling routine that infers the number of molecules that are active in the current segment. To that end, robust descriptors are derived for each segment: average photon counts, SBRs, and the visibility/contrast of the emission signal. These features are used to identify high-confidence examples of background, single- and multi-fluorophore segments. A deliberately conservative approach is taken: only clear exemplars are labelled initially, and these seed labels are then propagated across the dataset with a k-means-based propagation algorithm. Outliers and ambiguous cases are aggressively excluded to maintain reliability of the assignment.

We then proceed to the calibration stage to obtain system parameters, such as the initial visibility of the illumination minimum $\nu_0$ or expected single-molecule brightness and background levels. These values act as priors for subsequent model fits. Only traces with sufficient contrast are used for calibration, ensuring that the priors are robust.

Within blocks containing active emitters, a motility analysis is performed to capture dynamical behavior of the origami. Groups of contiguous blocks are examined for signatures of motion of the COM, and when present, a HMM is fitted to the time series of positions. In fact, we fit 10 randomly initialized HMMs for each number of states (ranging between 1 and 5) in order to remain agnostic about the apparent dynamics of the system. For each trace, the best model is selected according to a Bayesian information criterion. This yields per-tuple assignments of latent motility states together with statistics of the binding events, allowing quantification of dynamic transitions.

Last, we co-localize the two molecules with a MLE. The trace is bootstrapped into overlapping chunks, either according to a number of photons per bin or a time interval for the chunks and a specified stride to move the bin. Each chunk is fitted to a parametric PSF model under a Poisson likelihood, stabilized by the calibration priors. Overlapping chunks both increase temporal resolution and provide bootstrap-like uncertainty estimates.

After having obtained time-resolved emitter positions, we resolve ambiguities arising from the ill-behaving likelihood. Since the measurement remains ambiguous with respect to permutations and mirroring of the obtained positions, we sort and mirror the positions as to minimize their overall pair-wise distance.

In summary, the workflow proceeds as follows:

1. compress and segment raw traces at bleaching steps,
2. label segments conservatively with number of active molecules,
3. detect dynamical states and dwell times,

locally calibrate PSF-parameters and background against trusted single-molecule references, i.e. calibrate per trace if possible,infer positions of two molecules simultaneously with MLE.

#### Derivation of COM localization uncertainty and brightness

In order to derive the center-of-mass localization uncertainty for two fluorescent molecules with finite separation $d$ and COM $x_0$, we consider a quadratic intensity profile of the form $y_i = a\,x_i^2$ to probe the markers at three distinct positions $x_i = \pm\frac{L}{2}, 0$. The expected response of the two markers reads

$$y_i = a\left(x_i - \left(x_0 - \frac{d}{2}\right)\right)^2 + a\left(x_i - \left(x_0 + \frac{d}{2}\right)\right)^2 + b,$$

where a denotes the normalization (brightness) of the incident beam and b is a background parameter. The mean intensity $y_i$ of the i-th exposure can be simplified to

$$y_i = a\left(2(x_i - x_0)^2 + \frac{d^2}{2}\right) + b,$$

and the derivative with respect to $x_0$ is given by $\partial_{x_0} y_i = 4a(x_0 - x_i)$. From that, we can calculate the Fisher Information J with respect to the COM $x_0$ for a Poisson variable, near $x_0 = 0$:

$$J_{x_0} = \sum_i \frac{\left(\partial_{x_0} y_i\right)^2}{y_i} = \frac{8a^2L^2}{b + a\frac{L^2 + d^2}{2}}.$$

Since the localization uncertainty (Cramer Rao Bound) of $x_0$ is given by $\sigma_{x_0} = 1/\sqrt{J}$, we can calculate the ratio of the localization uncertainties $\sigma_{x_0}(d \neq 0)/\sigma_{x_0}(d = 0)$ from the ratio of Fisher Informations

$$\frac{\sigma_{x_0}(d \neq 0)}{\sigma_{x_0}(d = 0)} = \sqrt{\frac{J_{x_0}(d = 0)}{J_{x_0}(d \neq 0)}}$$

This readily yields

$$\frac{J_{x_0}(d = 0)}{J_{x_0}(d \neq 0)} = 1 + \frac{d^2}{L^2}\frac{SBR}{SBR + \frac{2}{3}}.$$

Similarly, we calculate the ratio of the expected mean intensity among the three exposures for either $d = 0$ and finite separation, which yields:

$$\frac{I_{fluor}(d \neq 0)}{I_{fluor}(d = 0)} = 1 + \frac{3d^2}{2L^2}\frac{SBR}{SBR + 1}.$$

### Figures

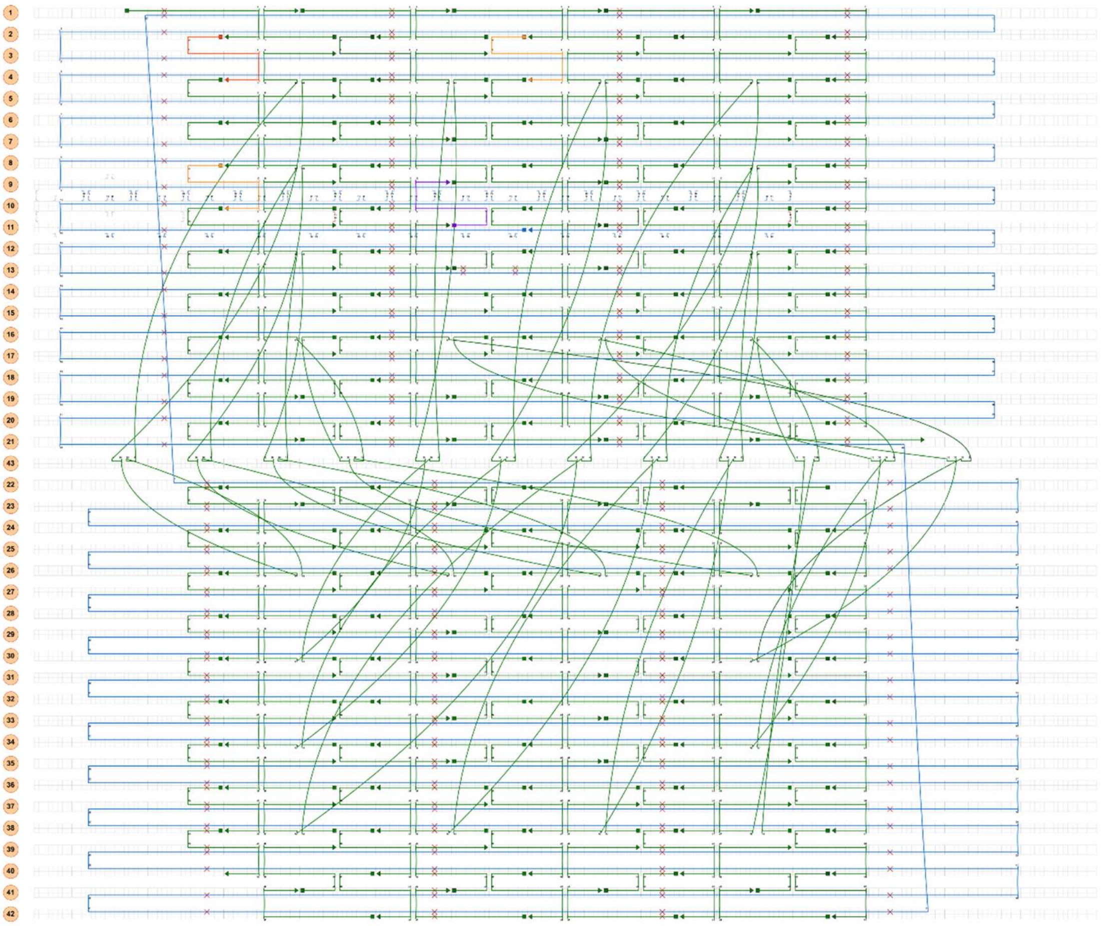

**Fig. S1: Cadnano design of the DNA origami.** The p8064 scaffold strand is shown in blue. Orange staples are extended at the 5′ end by two thymidine spacers followed by seven unpaired nucleotides (5′-AGTCCTA-3′), forming transient binding sites. The red staple is extended at the 5′ end by four thymidine spacers followed by an Atto 647N dye. The purple staple is extended at the 3′ end by three thymidine spacers and 15 nucleotides that form the base of the hinge. The design was adapted from (1,2).

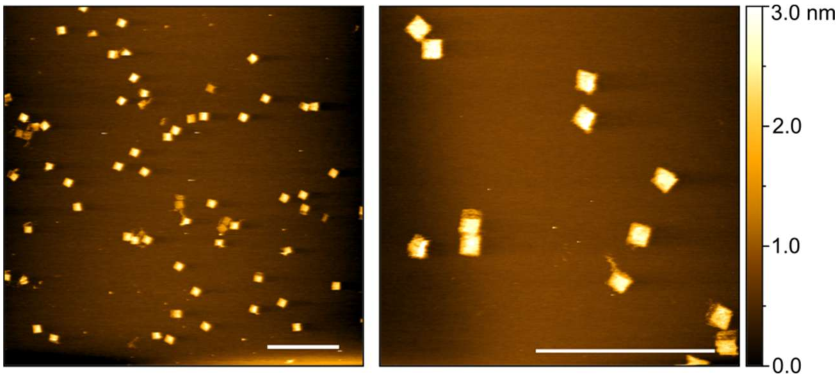


**Fig. S2: Atomic force microscopy images of annealed and purified DNA origami structures.** Scale bars: 500 nm.

#### Tables

**Table S1: Staple strands used to assemble the DNA origami sheet.** Staple strand positions are given as B_H5′[p5′]_H3′[p3′], where the first helix number and bracketed index specify the staple's 5′ start (helix number and position within that helix), and the second pair specifies the staple's 3′ end. Sequences are listed in 5' to 3' direction.

| Position | Sequence |
|---|---|
| B_41[56]_39[63] | TATCATTCAAATAAACAGCCATATAGAATAACATAAAAAC |
| B_21[152]_19[151] | GAGAATCGCTGAGAGACTACCTTTCGCTATTA |
| B_2[127]_1[119] | GTCTGGCCTTCCTGTTCATATGT |
| B_32[135]_34[136] | CCTCATAACCCTCAGAACCGCCTAATAAGT |
| B_40[167]_42[168] | CTTAAATCGCAAATCAGATATAGAAGGCATTT |
| B_9[120]_7[119] | AATTGCGGCGGGCCGTTTTCACTGCTCGTC |
| B_28[63]_27[63] | AAATTGGGCTTGAGATGACGTTGGGAAGAAAA |
| B_36[63]_35[63] | CCTTATTAGCGTTTGCAGAACCGCCACCCTCA |
| B_16[95]_29[159] | TATCTTTATTTGGAACGAGCAGACGGTCAATCATA |
| B_16[159]_37[159] | CAATCAATTTTCAGAAGGAAAGGTGAATTATCACC |
| B_2[63]_1[55] | ATTCTCCGTGGGAACATGCCTGAG |
| B_32[167]_34[168] | AACTACAACCCTCAGAGCCACCACACATGGCT |
| B_38[63]_3[159] | AAACGTAGTTTTTTCCCAGGGCTGCGCAACTGTTG |
| B_23[56]_22[40] | TAGTAGTAAAATTAATGCCGGAGAGGGTAGCT |
| B_4[71]_6[72] | CACCGGAAAAACTTAAATTTCTGGCAAGAA |
| B_38[39]_40[40] | GGTGGCAGCAGCCTTTACAGAGTATTTATC |
| B_28[159]_27[159] | CAAGAACCGGATATTCACATTCAACTAATGCA |
| B_30[71]_32[72] | AATACGTAAAAGGAGCCTTTAATTGTATGGGA |
| B_1[152]_2[168] | ACAGGAAGATTGTATAAGCAAATATCAGCT |
| B_41[152]_39[159] | AATAGCAAAAGATTAGTTGCTATTGATAACCCACAAGAAT |
| B_22[167]_24[168] | AATGCAATAAACATTATGACCCTGAATATAAT |
| B_4[167]_6[168] | TGGGTAACTAGTGATGAAGGGTAGTCGCTG |
| B_30[39]_32[40] | AAAACGACACGTTGAAAATCTCTTCAGCGG |
| B_24[159]_23[151] | TCAACATGTTTTAAATAAATCGGT |
| B_21[88]_19[87] | TATCATATTGCAAATCCAATCGCATTTTAATG |
| B_20[71]_21[87] | GAACGCGAATTACTAGAAAAAGCCTGTTTAG |
| B_1[19]_2[40] | TTTTGATCTACAAAGGCTATCAGGTCATAACGGCGG |
| B_32[63]_31[63] | AACAACTTTCAACAGTCAAAAAAAAGGCTCCA |
| B_4[135]_6[136] | TTGTAAAAAATCCCGTAAAAAAAGTCATAACG |
| B_14[39]_16[40] | CATTCTGGACAATATTTTTGAATGAATACATT |
| B_8[103]_10[104] | CGGGGGTTCTAATGAGTGAGCTAATGAGAGAG |
| B_4[39]_6[40] | ATGTTTACTCACGGAAAAAGAGACTGCCATCC |
| B_36[95]_35[87] | GCGTTTTCATCGGCAAGCCGCC |
| B_23[120]_22[104] | ATAAAGCCGGGTGAGAAAGGCCGGAGACAGTC |
| B_38[95]_7[159] | ATTAAGACTTTCTGTGGTGGTTCAGCAAATCGTTA |

| | |
|---|---|
| B_11[120]_9[119] | CGTGAACTTCACCGCCTGGCCCCTCACATT |
| B_34[167]_36[168] | TTTGATGAATTCACAAACAAATAAGTCACCAA |
| B_36[39]_38[40] | CATAATCTTGTCACAATCAATATACATAAA |
| B_16[127]_33[159] | GTTGAAATTTTGTACTGGACCCTCAGAACCGCCA |
| B_12[135]_14[136] | GGCGCTAGCATCACGCAAATTAACGAAAAACG |
| B_12[103]_14[104] | CGCTGCGCTCAGTGAGGCCACCGAACGCTCAA |
| B_16[167]_18[168] | CAAACCCTCTGATTGTTTGGATTTAAAACA |
| B_24[39]_26[40] | AATGGTCTGACTATTATAGTCAGCTTTAAA |
| B_2[71]_4[72] | CCCGTCGGCAGGAAGATCGCACTATAACCT |
| B_14[103]_16[104] | TCGTCTGATAAAAATACCGAACGAGGAAGGTT |
| B_34[135]_36[136] | TTTAACGGGTTGAGGCAGGTCAAATCAGTA |
| B_14[135]_16[136] | CTCATGGATAAAACAGAGGTGAGGAGTTGGCA |
| B_7[88]_5[95] | GCTGGTAACTGGTCAGCAGCAACCCTCATTTGCCGCCAGC |
| B_30[167]_32[168] | AGCGAAAGACCATCGCCCACGCATCCAGTACA |
| B_13[88]_11[87] | TTTTATAAGTAACCACCACACCCGCAAAGGGC |
| B_38[71]_40[72] | ATGTTAGCAGGGAAGCGCATTAGATAATTTGC |
| B_32[159]_31[159] | CGCCTGTAGCATTCCAGCCGACAATGACAACA |
| B_30[159]_15[95] | ACAGCATCTTTGGAGCACTCCTAAAACATCGCCAT |
| B_21[120]_19[119] | TAAAGCCGTTGGGTTATATAACGTGAGTGA |
| B_34[39]_36[40] | TTAAGAGAGAGCCGCCACCCTCCATCTTTT |
| B_10[39]_12[40] | GAAATCGGCCAGTTTGGAACAAGACTATGGTT |
| B_16[63]_25[159] | AGAGCCGTTTTGCAAAAGAAAGCAAACTCCAACAG |
| B_40[71]_42[72] | CAGTTACACAAGAACGGGTATTAAAATGCAGA |
| B_26[39]_28[40] | CAGTTCACATTATACCAGTCAGGGTTTAAT |
| B_8[167]_10[168] | TGCGCGCCCCTGTCGTGCCAGCTGGGTGGT |
| B_12[159]_37[127] | GAAGGGAATTTGGAATACCGAAGGTAAATATTGAC |
| B_4[103]_6[104] | GAGGTGGAAGTTGGGCGGTTGTGTTCAGCGTG |
| B_22[135]_24[136] | TTCAAAATCAGAGCATAAAGCTATGCAACT |
| B_29[88]_27[95] | ATAAATTGTGCCCTGACGAGAAACGAACTAACGGAACA |
| B_36[127]_35[119] | AATCAAGTTTGCCTTTGCCAGCAT |
| B_20[39]_21[55] | GTTAATTTGGCGTTAAATAAGAATAAACACCG |
| B_18[167]_20[168] | GAAATAAATCCCTTAGAATCCTTATCAAAA |
| B_28[167]_30[168] | AATCTTGAAGGGAACCGAACTGACCCCTCAGC |
| B_14[95]_13[87] | AATGGATTATTTACATAGAAGTGT |
| B_24[127]_23[119] | GGTGTCTGGAAGTTTCATTAAGCA |
| B_42[71]_41[55] | ACGCGCCTGTTTATCAACAATAGATCTTTCCT |
| B_4[159]_37[63] | GCCAGGGTTTTAAAATACAGAAAATTCATATGGTT |
| B_30[63]_3[95] | ATGCCACTTTTGGGAACGGCCAGCCAGCTTTCCGG |
| B_24[63]_23[55] | TTTGACCATTAGATACTCTACTAA |
| B_26[95]_7[63] | CCAATACTTTTTCCTCACAGCGCGGTTGCGGTATG |

| | |
|---|---|
| B_35[88]_33[87] | ACCAGAACAACAGTTAATGCCCCTAGCCCG |
| B_32[39]_34[40] | AGTGAGAGCTCAGTACCAGGCGTGAAAGTA |
| B_41[88]_39[95] | ACCGCACTCGTCTTTCCAGAGCCCGGGAGAATTAACTG |
| B_9[88]_7[87] | TGGGGTGCTCTGCCAGCACGCGTGGCCCTGCG |
| B_14[71]_16[72] | TTCACCAGGCGCGAACTGATAGCAACAACT |
| B_4[127]_33[63] | CGACGGCTTTCTGAAACAGATAAGTGCCGTCGAG |
| B_31[120]_29[119] | AACAGCTTAGAGGCTTTGAGGACTCTGCTCCA |
| B_22[71]_24[72] | TAGCTGATGCATTAACATCCAATAGAACGAGT |
| B_12[167]_14[168] | CGAGAAAGCTTTGATTAGTAATAAGAACAA |
| B_4[95]_29[63] | GCCGCCACTTTACGAAGGCCGCGAAACAAAGTACA |
| B_14[63]_13[63] | TCACACGACCAGTAATTTTTAGACAGGAACGG |
| B_19[120]_17[127] | ATAACCTATGAATATACAGTAATCATCATATTCCTGA |
| B_2[135]_4[136] | TAATTCGCCGCCATTCGCCATTCATCACGACG |
| B_36[71]_38[72] | CATAGCCCTACCAGCGCCAAAGACTACGCAGT |
| B_33[120]_31[119] | GTACCGCCGTTAGCGTAACGATCTATTTCTTA |
| B_36[135]_38[136] | GCGACAGGGAAATTATTCATTAAACCGAGG |
| B_26[71]_28[72] | ATATTCATATCTACGTTAATAAAACACCAGAA |
| B_26[167]_28[168] | ATAGCGAGGATACATAACGCCAAATCAAGAGT |
| B_6[39]_8[40] | CACGCAACGCACTCAATCCGCCGGGTTGAGGA |
| B_35[120]_33[119] | TGACAGGAGGGTCAGTGCCTTGAGGAGGTTTA |
| B_38[127]_11[159] | ATAATAACTTTGAAAGCGATTTTTTGGGGTCGAGG |
| B_16[71]_18[72] | AATAGATTCATTATCATTTTGCGGATTCGC |
| B_42[135]_41[119] | ACAAAAGGTAAAGTAATTCTGTCGCCGTTTT |
| B_42[167]_41[151] | TCGAGCCAGTAATAAGAGAATATACCGCGCCC |
| B_8[159]_37[95] | TGTGCACTTTTTCCTTATAAAAGGGCGACATTC |
| B_36[167]_38[168] | TGAAACCAGTCACCGACTTGAGCCAAGCAGAT |
| B_18[71]_20[72] | CTGATTGCCATTTGAATTACCTTAGACAAA |
| B_24[167]_26[168] | GCTGTAGCGTCAGGATTAGAGAGTAAACCAAA |
| B_36[159]_35[159] | TCGATAGCAGCACCGTGACGATTGGCCTTGAT |
| B_22[39]_24[40] | ATTTTTGAAAGGTGGCATCAATATTTCGCA |
| B_10[167]_12[168] | TTTTCTTTTGCCGTAAAGCACTAAACGTGG |
| B_18[135]_20[136] | AACGTCAGTGCTTCTGTAAATCGTTTAACCTC |
| B_20[135]_21[151] | CGGCTTAGAACGCTCAACAGTAGGGCTTAATT |
| B_24[95]_23[87] | GATTCCCAATTCTGCAATCATA |
| B_34[159]_15[127] | TACAGGAGTTTGGAATTGAACCACCAGCAGAAGA |
| B_4[63]_25[63] | ACAATCGGTTTCCTCAAATGAAGCAAAGCGGATTG |
| B_10[63]_9[63] | AAAATCCTGTTTGATGCCACACAACATACGAG |
| B_8[63]_25[95] | CGTGAGCCTTTGCGGAATAGGAAGCCCGAAAGA |
| B_10[159]_9[159] | TCACCAGTGAGACGGGCTTTCCAGTCGGGAAA |
| B_8[135]_10[136] | AGAATGCGTTGCGCTCACTGCCCGCAACAGCT |

| | |
|---|---|
| B_30[135]_32[136] | CGGCTACGATACCGATAGTTGCCAGACAGC |
| B_2[159]_1[151] | AACCAATAGGAACGCCCCCCAAAA |
| B_12[63]_25[127] | CGCTACAGTTTTAAAATGTTTAATTCGAGCTTCAA |
| B_40[135]_42[136] | AGCTACATCGTAGGAATCATTAAAGTACCG |
| B_12[39]_14[40] | GCTTTGACAGGCCGATTAAAGGGAAAAAGGGA |
| B_7[120]_5[127] | ATAAACACGGACTTGTAGAACGACATCGACATAAAAA |
| B_14[127]_13[119] | AATACCTACATTTTGGTAAAAGA |
| B_14[159]_13[159] | CCAGCCATTGCAACAGCGTTGTAGCAATACTT |
| B_26[127]_11[63] | GGTAATAGTTTGGCGCGTAGTCCACTATTAAAGAA |
| B_31[88]_29[87] | TTTATCAGGGAAGTTTCCATTAATCGCCTG |
| B_10[135]_12[136] | GATTGCCCCATCACCCAAATCAAGAAGGAGCG |
| B_6[135]_8[136] | GAACGTGCTCCCTTACACTGGTGTCTGCGGCC |
| B_28[39]_30[40] | TTCAACTAGCGATTATACCAAGACCAACCT |
| B_14[167]_16[168] | TATTACCGCGCCTGCAACAGTGCCAAATAT |
| B_26[63]_3[63] | TGAATCCCTTTCGAAACGTACGACAGTATCGGCCT |
| B_26[159]_15[63] | AGGCTTTTTTTCAATAGATGCTATTAGTCTTTAAT |
| B_23[152]_22[136] | TGTACCAAGCCTGAGTAATGTGTAGGTAAAGA |
| B_28[71]_30[72] | CGAGTAGTACGGAGATTTGTATCAACGGGTAA |
| B_16[135]_18[136] | AATCAACATTATCAGATGATGGCATCAGGTTT |
| B_12[71]_14[72] | TAATGCGCTACGCCAGAATCCTGTGGCAGA |
| B_33[88]_31[87] | GAATAGGTGTAAATGAATTTTCTGTATCGG |
| B_24[135]_26[136] | AAAGTACAGCGAACCAGACCGGAGTTTTGC |
| B_18[39]_20[40] | CGCAGAGGAAGAAAACAAAATTAAATATTTTA |
| B_2[167]_4[168] | CATTTTTTGGAAGGGCGATCGGTATTAAGT |
| B_38[159]_15[159] | AAAGTTACTTTATCTGGTCCGGTCAGTATTAACAC |
| B_23[88]_22[72] | CAGGCAAGATCAATATGATATTCAACCGTTC |
| B_42[103]_41[87] | GACAATAAACAACATGTTCAGCTACCAAGT |
| B_20[167]_21[187] | TCATAGGTCCATATTTAACAACGCCAACATGTTTT |
| B_26[135]_28[136] | CAGAGGGAGATTTAGGAATACCATTACCCA |
| B_20[103]_21[119] | AATGCTGAGCGTTATACAAATTCTTACCAGTA |
| B_29[120]_27[127] | TGTTACTTGTAACAAAGCTGCTCAAAAGATTCATCAGTTG |
| B_6[103]_8[104] | GTGCTGGTTGGGTAAAGGTTTCTTGGTCATAC |
| B_40[103]_42[104] | CTAACGAGCATCGAGAACAAGCAACAGACGAC |
| B_38[167]_40[168] | AGCCGAACTGAGTTAAGCCCAATATTTGAAGC |
| B_16[103]_18[104] | ATCTAAAAAGAAGGAGCGGAATTACAGTACCT |
| B_18[103]_20[104] | TTTACATCACATAAATCAATATATTATATGTA |
| B_24[71]_26[72] | AGATTTAGCATCAAAAAGATTAAGCGTCATAA |
| B_13[120]_11[119] | GTCTGTCGGCGCTGGCAAGTGTGCCCACTA |
| B_6[167]_8[168] | GCAGCCTCACGGCATCAGATGCCGTGTCAC |
| B_1[56]_2[72] | AGTCTGGAGCAAACAAGAGAATCGTAACAA |

| | |
|---|---|
| B_28[135]_30[136] | AATCAACAGCCGGAACGAGGCGGGTAGCAA |
| B_10[71]_12[72] | AGCAGGCGCGTGGACTCCAACGTCCGCGCT |
| B_38[135]_40[136] | AAACGCACGCTAATATCAGAGATTGCACCC |
| B_19[88]_17[95] | GAAACAGTGGGAGAAACAATAACGGAACAAAGAAACCACC |
| B_1[88]_2[104] | GTAATCGTAAAACTAGCATGTCAAAGCCAGCT |
| B_16[39]_18[40] | TGAGGATTCCGAACGTTATTAATTCAAAATCG |
| B_2[95]_1[87] | CATTAAATGTGAGCGAGATGAACG |
| B_21[56]_19[55] | GAATCATAGAAAACTTTTTCAAATTTACATTT |
| B_34[71]_36[72] | TTCGGAACGAGCCACCACCCTCAGTTTTCGGT |
| B_32[71]_34[72] | TTTTGCTAAGGGTTGATATAAGTACTGCCTAT |
| B_34[63]_3[127] | CTATTATTTTTCAGTGCCAGGAAACCAGGCAAAG |
| B_8[71]_10[72] | TTCGCGTCCCGGAAGCATAAAGTTTGCCCC |
| B_41[120]_39[127] | TATTTTCAATTTTATCCTGAATCTTCAGAGGGTAATTGAG |
| B_10[103]_12[104] | TTGCAGCAGTCTATCAGGGCGATGAGCGGTCA |
| B_1[120]_2[136] | ACCCCGGTTGATAATCAGAAAAGATCAAAAA |
| B_6[71]_8[72] | TGCCAACGAGCCGGGTCACTGTTCCTGTTC |
| B_6[159]_5[159] | CGGCCAGAGCACATCCCCGCACAGGCGGCCTT |
| B_19[56]_17[63] | AACAATTTTTTGAATACCAAGTTATTAAAAGTTTGAGTAA |
| B_19[152]_17[159] | ATTAATTTGAAATTGCGTAGATTTATTCATCAATATAATC |
| B_6[63]_5[63] | GCAGCACCGTCGGTGGGCAGAAACAGCGGATC |
| B_30[103]_32[104] | TTTCATGACTTGCTTTCGAGGTGAAAAGTTTT |
| B_22[103]_24[104] | AAATCACCGCAAAGAATTAGCAAAATTCCATA |
| B_28[103]_30[104] | ATAAGGCTTGTCGAAATCCGCGACAAAGACTT |
| B_32[103]_34[104] | GTCGTCTTTCCAGACGTTAGTATCACCGTACTCAGTAACAGTG |
| B_36[103]_38[104] | ACTGTAGCAACCGATTGAGGGAGGCAAAAGAA |
| B_38[103]_40[104] | CTGGCATGAACACCCTGAACAAAGTACCAACG |
| B_26[103]_28[104] | GATAGCGTACATTATTACAGGTAGTTCAGTGA |
| B_24[103]_26[104] | TAACAGTTCTTCAAATATCGCGTTTTAGACTG |
| B_34[103]_36[104] | CCCGTATACACCACCAGAGCCGCCAGCGTCAG |

**Table S2: Modified strands used in the DNA origami.** Staple strand positions are given as B_H5′[p5′]_H3′[p3′], where the first helix number and bracketed index specify the staple's 5′ start (helix number and position within that helix), and the second pair specifies the staple's 3′ end. Sequences are listed in the 5′ to 3′ direction. Regions corresponding to the main double-layer structure are shown in black; thymidine spacers in yellow; transient binding regions in red; hinge sequences in blue; and fluorophore modifications in dark red.

| Position | Name | Sequence |
|---|---|---|
| B_2[103]_4[104] | Binding_site_1 | AGTCCTATTTTCATCAACACCGCTTCTGGTGCCAGCTTTCA |
| B_8[39]_10[40] | Binding_site_2 | AGTCCTATTTCCCCGGGTTATCCGCTCACAATTGTGGTTCC |
| B_2[39]_4[40] | Static_fluorophore | /Atto647N/TTTTATTGACCGCCAGTTTGAGGGGACGACAGCGCC |
| B_11[88]_9[87] | Hinge_base | GAAAAACCAGCGGTCCACGCTGGTGTAAAGCCTTTGACACTTCGTGTGTG |
| Hinge | Hinge_binding_site | CGTGGATCACACGAAGTATCACACACGAAGTGTTTAGGACT |
| Hinge | Mobile_fluorophore | /Atto647N/TACACTTCGTGTGTGATACTTCGTGTGATCCACGCACACACGAAGTGTC |

**Movie S1: Nonstop tracking of two fluorophores at the nanoscale.** Animation of the co-tracking of two randomly moving fluorophores at the nanometer scale. The latest positions of the two fluorophores are indicated by the yellow stars. Heatmaps show the aggregated localizations and reveal the underlying structure of the DNA construct with it’s three states: two bound states and one unbound, freely-diffusing state.
