## Supplementary figures and images for "Nonstop nanometric resolution of randomly moving point scatterers with focused light"

### Movie S1

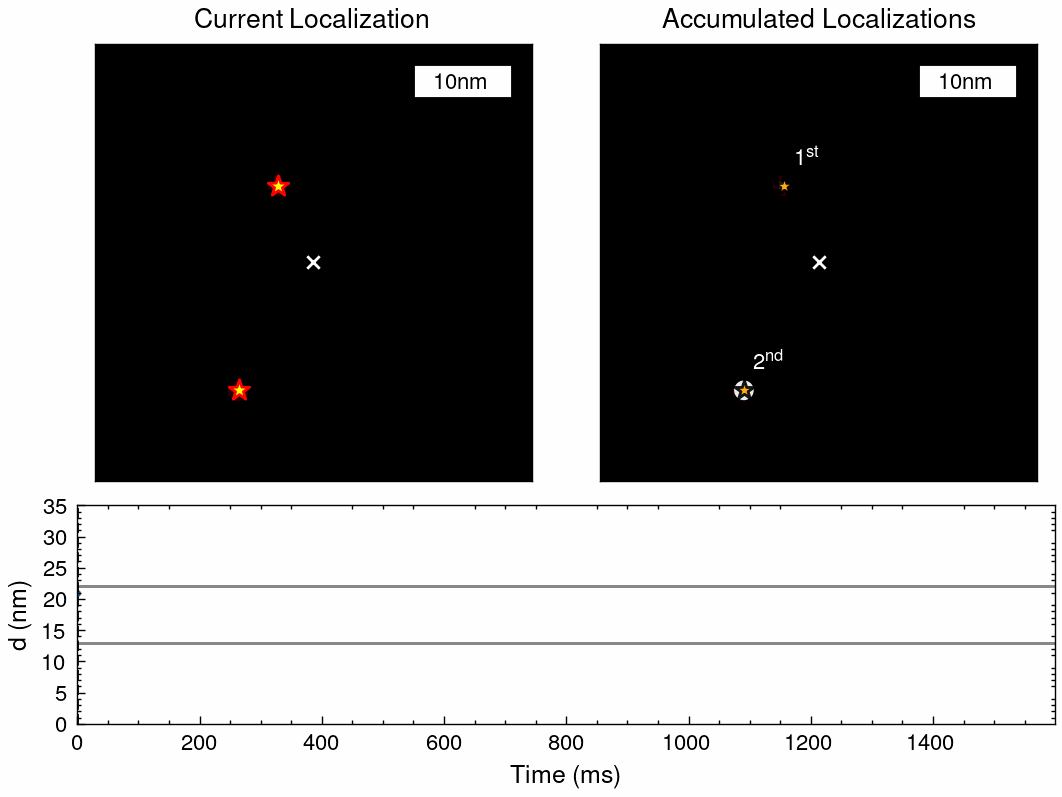
